# Dynamic behavioral and molecular changes induced by chronic stress exposure in mice

**DOI:** 10.1101/2021.05.07.443011

**Authors:** Thomas D. Prevot, Dipashree Chatterjee, Jaime Knoch, Sierra Codeluppi, Keith A. Misquitta, Corey J.E. Fee, Dwight Newton, Hyunjung Oh, Etienne Sibille, Mounira Banasr

## Abstract

Depression is a leading cause of disabilities around the world, and the underlying mechanisms involved in its pathophysiology are broad and complex. Exposure to chronic stress is a risk factor for developing depressive-symptoms and contributes to cellular and molecular changes precipitating the emergence of symptoms. In the brain, excitatory neurons, inhibitory interneurons and supporting astroglial cells are all sensitive to chronic stress exposure and are known to be impaired in depression.

Using an animal model of chronic stress, we assessed the impact of variable durations of chronic stress on the emergence of behavioral deficits and associated molecular changes in the prefrontal cortex (PFC), brain region highly sensitive to stress and impaired in depression. Mice were exposed to up to 35 days of chronic restraint stress and were assessed weekly on behavioral tests measuring anxiety and anhedonia. PFC Protein and RNA levels of specific markers of excitatory, inhibitory synapses and astroglia were quantified using western blot and qPCR, respectively. Correlation and integrative network analyses were used to investigated the impact of chronic stress on the different compartments.

Results showed that chronic stress induces anxiety-like behaviors within 7 days, while anhedonia-like behaviors were observed only after 35 days. At the molecular level, alterations of many markers were observed, in particular with longer exposure to chronic stress. Finally, correlation analyses and integrative network analyses revealed that male and female mice react differently to chronic stress exposure and that some markers seem to be more correlated to behaviors deficits in males than in females.

Our study demonstrate that chronic induces a dynamic changes that can be observed at the behavioral and molecular levels, and that male and female mice, while exhibiting similar symptoms, have different underlying pathologies.

## Introduction

Major depressive disorder (MDD) is one of the most prominent psychiatric illnesses worldwide affecting 322 million people worldwide [1] and causes an enormous economic strain [2]. MDD symptoms include anhedonia and feelings of worthlessness/helplessness [3], and is highly comorbid with anxiety disorders (^~^50-60%) [4, 5]. It is also a multifaceted disease with various contributing risk factors including, genetics, sex, and environmental stress [6]. It has been reported that MDD is more prevalent in women than men [7], with women experiencing more severe forms of MDD [8, 9], and being most susceptible to stress [10]. Although the scope and knowledge behind psychiatric therapy has greatly improved, increasing evidence has shown that the efficacy of current antidepressants is limited. Indeed, only ^~^60% of patients respond effectively to treatment [11] and others often see severe side effects and blunting effects [12]. This lack of efficacy is mostly due to the fact that current antidepressants do not specifically target the primary underlying cellular pathologies of MDD [12]. This is not surprising considering the origins of these drugs are largely attributed to serendipity [12, 13], and that rely mostly on compounds targeting the monoaminergic deficiencies, which are not effective in all cases [12]. Taken together, the heavy burden on society and individuals and its insufficient pharmacological treatment highlights the critical need for a better understanding of the primary pathophysiology of MDD, as a first step toward developing new therapeutic approaches.

In recent years, growing evidence suggests that MDD is characterized by morphological pathologies where brain regions, cells and cell compartments lose integrity and function [14]. Evidence from human post-mortem and preclinical studies associates structural and cellular abnormalities in corticolimbic brain regions including the prefrontal cortex (PFC) with the pathophysiology of the MDD [15–17]. Changes reported implicate dysfunctions within 3 main compartments of the neural circuitry: excitatory synapses, GABA interneurons, and astroglial cells. Indeed, postmortem and preclinical studies consistently report synaptic loss [18–21], GABAergic dysfunction [22–26], and astroglial abnormalities [27–31] in the PFC of brains from patients with MDD, or from animal models of chronic stress. More precisely, reduction in key markers of GABA neurons such as GAD67, paravalbumin, VIP somatostatin (SST))[32, 33] and of astroglial including GFAP and GS [28–31, 34] were reported in MDD. Interestingly, despite their intricate functional roles, the cellular alterations occurring within the GABAergic, astroglial and synaptic (i.e. pyramidal neurons) compartments of the tripartite synapse in MDD are usually studied independently when the illness is already fully developed, preventing any understanding of their gradual emergence.

Chronic exposure to stress is a major risk factor of MDD and is an ideal tool to be used in preclinical models to better understand the potential changes leading from chronic stress state to MDD pathology [35]. Studies showed that females are also more sensitive to stress exposure [10, 36], and tend to develop more severe symptoms than male counterparts [8, 9]. In the PFC, it is commonly reported that chronic stress induces synaptic loss, in number and/or reduced presynaptic (Syn1 or VGLT1) and presynaptic (PSD95 and gephrin) marker expression. Some of these changes are shown to contribute to behavioral deficits [36, 37]. GABAergic [10, 26] and astroglial functions [36, 38] are also altered in the PFC mice or rats subjected to chronic stress.

In preclinical settings, studies consistently utilize chronic stress exposure to induce behavioral and cellular changes in rodents similar to what can be seen in depressed patients [4, 39–41]. Several models are used to achieve chronic stress state including exposure to unpredictable chronic mild stress, chronic restraint stress (CRS), foot shock, social isolation, and social defeat [42]. Here, we opted for the use of CRS as studies showed it is reliable in inducing depressive-like behaviours, cellular and morphological changes in neuronal morphology in the PFC [19, 42–44].

Here, we investigated if chronic stress alters expression of the GABAergic, glutamatergic and astroglial markers differentially throughout the chronic stress exposure. We hypothesized that the trajectory of behavioral changes may map onto the compartment-specific dysfunction during chronic stress exposure. We also used co-expression analyses to investigate the relationship between markers in order to determine how exposure to chronic stress shifts expression profiles, interestingly revealing differential progression between males and females.

## Materials and Methods

### Animals

8 week-old C57BL/6 mice from Jackson Laboratories (Stock No: 000664; Bar Harbor, Maine, USA) were used. Prior to experimentation, animals were habituated to the facility for 1 week with a 12h light/dark cycle and *ad libitum* access to food and water. The animals were assigned either the control group or a chronic restraint stress (CRS) animals subjected to 7, 14, 21, 28 or 35 days of CRS (n=16/group, 50% females). The control group was handled daily for 3 days and given nesting material [45]. Control mice were housed in a separate room than the CRS groups to avoid any disruption. All experiments were conducted in line with guidelines provided by the Canadian Animal Care Committee and protocol was approved by the animal care committee of the *Centre for Addiction and Mental Health*.

**Chronic Restraint Stress** (CRS) protocol consists of placing the mice in a Falcon^®^ Tube, with a hole at the bottom and on the cap to allow air to flow through [46]. Mice stayed in the tube for 1hr before being released, twice a day (2 hours apart), every day, for the duration of time they were to be subjected to CRS (7, 14, 21, 28 or 35 days).

**Coat State and Body Weight** were measured weekly. Coat state was assessed following the method described by Yalcin et al.[47]. Weight gain, as a percentage, was measured using the weight of each mouse on Week 0 as the reference.

**Sucrose Consumption** was performed weekly. Mice underwent a 48hr habituation period to a 1% sucrose solution (only for the first exposure – subsequent habituation only lasted 24hrs). Then the mice were fluid deprived for a 16hr overnight (^~^6pm to 10am). On the subsequent morning, the sucrose solution was returned to the mice for a 1hr period after which consumption was recorded. This same protocol is repeated as a measure water consumption to be used for comparison. Each week the ratio of consumption was analyzed as ratio of consumption which is calculated as: 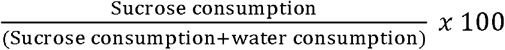.

### PhenoTyper

The PhenoTyper test was used on a weekly basis to assess anxiety-like behavior as per the protocol described in Prevot et al., 2019. [44] (see supplementary methods for detailed description of the assay). Briefly, mice are placed in the PhenoTyper^®^ boxes overnight, in which their overall activity is monitored by an infrared camera, mounted in the ceiling of the box, and linked to Ethovision^®^ software. At 11pm, and for 1hr, a light challenge was applied above the food zone, and time spent by the animal in the food zone (FZ) or the shelter zone (SZ) was monitored. Using the time spent in the FZ and SZ, two residual avoidance (RA) scores were calculated (one for each zone). This RA calculation informs us about the reaction after the mice after light challenge as compared to controls. Typically, control animals have a RA score at 0 while animals subjected to stress have consistently shown a positive RA score. This would suggest that animals undergoing stress actively avoid the zone where the light was shined even after it is turn off when they are displaying anxiety-like symptoms.

### Sample collection and preparation

Twenty-four hours after the last chronic stress session, brains were collected and PFC was dissected and frozen using on dry ice. Using a Qiagen Allprep RNA/protein Kit (#80404), RNA and proteins were extracted from the PFC samples. Isolated RNA was converted to cDNA using a SuperScript VILO cDNA Synthesis Kit (ThermoFisher, Massachusetts, USA, Cat#: 11754050). For the protein level in each PFC sample, a Pierce BCA (Bicinchoninic Acid) Protein Assay Kit (Thermofisher, Massachusetts, USA, Cat #: 23250) was used to measure protein concentration.

**Western Blot and qPCR** were performed as per standard in the field, and are fully described in **supplementary methods**. In brief, western blot experiments were carried on BioRad Criterion TGX Stain-Free Precast gels (4-20%), transferred on nitrocellulose membrane, incubated with antibodies (see **Supplementary Table 1**), enhanced luminescence with ECL, imaged and quantified using a molecular imager (ChemiDoc XRS, BioRad) with ImageLab™ software. qPCR were performed to amplify cDNA of somatostatin (SST), parvalbumin (PV) and vasopressin (VIP), on a Mastercycler real-time PCR machine (Eppendorf, Hamburg, Germany). Results were normalized to three validated internal controls (actin, GAPDH, and cyclophilin G) and calculated as the geometric mean of threshold cycles. All primer sequences can be found in **supplementary methods**.

### Statistical Analysis

Statistical analysis of behavioral outcomes and expression levels was conducted using StatView 5.0 software (Statistical Analysis System Institute Inc, NC, USA). Normal distribution of the data was assessed prior to statistical analysis. Data following a normal distribution were analyzed using ANOVA to determine main factor effects and followed by Dunnet’s post-hoc tests, when significant, in order to compare the impact of the different duration of CRS tested to the control group. Data with a non-normal distribution were analyzed using non-parametric Kruskal-Wallis test, followed by Bonferroni/Dunn post-hoc tests. Behavioral measurements, including coat state score, weight gain, residual avoidance in the shelter (SZ RA) and the food zone (FZ RA) as well as sucrose consumption were used to generate an overall z-score, as per the method described in Guilloux et al. [48]. Z-scores were calculated in a sex-dependent manner, using control mice of each sex as the reference group, with the formulae:

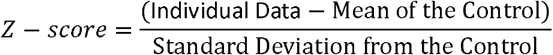

Each behavioral measure and z-scores were correlated to expression levels of the different markers. Pearson’s regression analyses were used for correlating markers with weight gain, FZ RA, SZ RA, sucrose consumption and z-score. Because of the ordinal distribution of the coat state scores, Spearman’s rank regression analyses were used for correlating markers with coat state scores. Markers expression levels were also correlated between each other using Pearson’s regression and FDR corrected for multiple comparison.

**Principal Component Analysis** (PCA) was performed taking into account expression levels from all markers, in R using the base “stats” package and visualized in 3 dimensions using the “plot_ly” package. The resultant loadings of each marker on the first 3 principal components were ploted with each PC. correlated with each PC.

### Network Analysis

See **Supplementary Methods** for details. Co-expression analysis was used to examine changes in coordinated expression between the markers as in [49]. All analyses were performed in R (version 3.6.0)[50]. Analytical code is available upon request. Briefly, Pearson correlation matrices between Z-normalized data, to account for marker scaling differences for all markers. Markers were then hierarchically clustered based on degree of correlation, and modules were generated from the resulting dendrogram using the dynamicTreeCutting function from the WGCNA R package [51], with a minimum module size of three. Networks were visualized in Cytoscape [52]. Co-expression modules were compared across stress groups using permutation testing to assess the degree to which they were preserved over the course of CRS. Differences in module composition in each CRS group versus controls, and between each CRS group were examined. Permutation testing (n=10,000) was used to compare cross tabulation-based measures of module preservation (i.e. whether markers remained co-expressed or not) and generate distributions of variability in module composition. These distributions were used to generate module-wise empirical p-values. Significant p-values represented a lesser degree of preservation (i.e. co-expression modules showed a different composition). Fisher’s p-value meta-analysis was used to combine module-wise p-values into a single p-value for each group-to-group comparison after Benjamini-Hochberg FDR correction [53].

## Results

### Chronic stress exposure induces a dynamic emergence of behavioral outcomes

In order to assess the impact of chronic stress in mice, weight gain, coat state scores and PhenoTyper test were performed every week for all animals (**Fig.1A**). Weekly measurement of weight gain and coat state are presented in **Supplementary Fig.1–2** and **Supplementary Results**. Between groups analyses were performed on Week 5, after completion of the different stress duration for each group. Kruskal-Wallis test (**Fig.1B**) identified significant difference in coat state between the groups (H=44.3, p<0.001). Bonferroni/Dunn *post hoc* test showed significant increase in coat state score with the increasing duration of the stress exposure (ps<0.05, for each CRS group, compared to control group). When sex was added as a co-factor no difference between groups was found.

**Figure 1:**
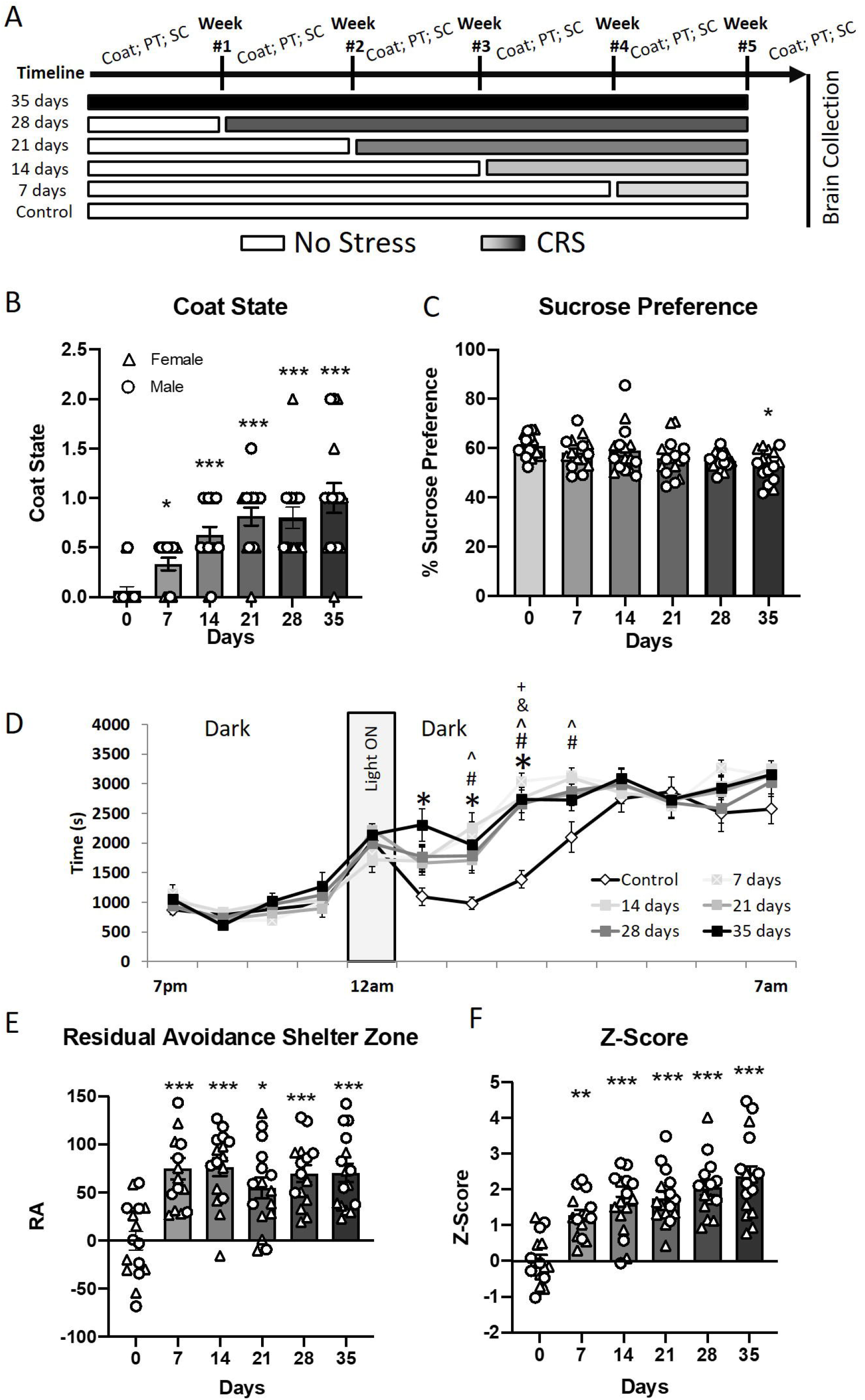
Chronic stress induces gradual emergence of behavioral changes. Mice (16/group; 50% females) were subjected to up to 35 days of chronic restrain stress (CRS). Their coat state score, anhedonia- and anxiety-like behaviors were assessed every week in the sucrose consumption (SC) and phenotyper (PT) tests, respectively (A). Coat state scores (B) showed gradual increase dependent of the duration of CRS exposure. Sucrose consumption (C) decreased gradually in mice subjected to CRS, with significant reduction compared to controls after 35 days of CRS. In the phenotyper test (D), mice were left overnight in the arena, where a light was shined from 11pm to 12am. Time spent in the shelter was measured, and we noticed that animals subjected to CRS spent more time in the shelter, even after the light challenge. Based on this observation, we calculated a residual avoidance (RA) score (E), which increased significantly in mice subject to CRS compared to controls. Finally, using a z-score approach, we see a gradual increase in z-score dependent of CRS duration, highlighting the gradual impact of CRS. *p<0.05, **p<0.01 and ***p<0.001 compared to controls (CRS0).

To detect CRS-induced anhedonia-like deficits, sucrose consumption was measured (**Fig.1C**). ANOVA revealed a significant effect of the CRS duration (F_(5,79)_=2.7; p=0.02) but no effect of sex or a duration*sex interaction. *Post hoc* Dunnet’s test identified a significant decrease in percent of sucrose consumed in the CRS35 group compared to control (p<0.05).

Animals were also tested weekly in the Phenotyper Test to assess anxiety-like behaviors throughout CRS exposure (**Supplementary Results and Supplementary Fig.3). Fig.1D** shows the time spent in the shelter zone (SZ) when the animals were tested on Week #5. Repeated measure ANOVA of the time spent in the SZ throughout the night shows a significant effect of CRS duration (F_(5;1080)_=4.39; p=0.0012), an effect of Sex (F_(1;1080)_=4.7; p=0.03), an effect of hour (F_(12;1080)_=150.2; p<0.001) but no duration*sex or duration*sex*hour Interaction (F_(5;1080)_=0.4; p>0.8). Dunnet’s test performed on the time spent in the shelter at each time point showed that there are no difference in time spent in the shelter between 7pm and 11am, and past 4am (**Supplementary Table 2**). However, Dunnet’s test showed that animals from the CRS35 group spent more time in the shelter than control at the 12am time point. This difference between the CRS35 group and the Control group was verified at the 1am, 2am and 3am time points. Significant differences compared to the control group were also observed for the CRS28 group at the 2am and 3am time point; for the CRS21 group at the 2am time point; for the CRS14 group at the 1am, 2am and 3am time points and for the CRS7 group at the 1am, 2am and 3am time points. In order to simplify the visualization of the differences observed between 12am and 4am, we used the RA calculation, as described in Prevot et al [44]. ANOVA performed on the RA score calculated from the time spent in the SZ revealed a significant effect of CRS duration (F_(5;82)_=10.23; p<0.0001) and sex (F_(1,82)_= 14.55; p<0.001) with no stress*sex interaction (**Fig.1E**). *Post hoc* analysis identified that animals subjected to CRS displayed greater RA-SZ compared to controls, for all CRS groups. The main effect of sex reflected that males from the CRS14, CRS28 and CRS35 groups exhibited an overall higher RA-SZ score than females (**Supplement Fig.4A**). Similarly to the analysis of RA score of the time spent in the SZ, the RA score of the time spent in the FZ was analyzed and showed similar results (**Supplement Fig.4B-C**).

Behavioral data were summarized using a z-score approach (**Fig.1F**). ANOVA of z-score showed a significant effect of CRS duration (F_(5,82)_=18.2, p<0.001), a significant effect of sex (F_(1,82)_=12.9, p=0.0006) and no CRS duration*sex interaction. *Post hoc* analysis revealed that in both males and females there is a significant increase of the z-score in all CRS duration groups, compared to the control group (ps<0.01). The effect of sex is explained by a Z-score higher in males than females (**Supplementary Fig.5**).

### Chronic stress exposure induces dynamic changes in cellular markers

Western blot analyses of astroglial (GFAP, GS, GLT1), GABAergic (GAD67, GPHN) and synaptic (vGLUT1, Syn1, PSD95) markers were performed on PFC samples obtained from all animals. ANOVA of GFAP protein expression levels, with sex as a cofactor, revealed a trend in main effect of CRS duration (F_(5;82)_=1.96; p=0.092), a significant main effect for sex (F_(1;82)_=7.59; p<0.01) and no CRS duration*sex interaction (**Fig.2A**). Dunnett’s *Post hoc* analysis identified a significant decrease in GFAP levels in the CRS28 group (p<0.01) when compared to controls. On this measure, males showed overall higher levels of GFAP than females (**Supplementary Fig.6A**). ANOVA of GS and GLT1 protein levels, with sex as a cofactor, revealed no main effect of CRS duration (F_(5:82)_=1.20; p=0.315, and F_(5:82)_=0. 9; p=0.44, respectively), sex, or CRS duration*sex interaction (**Fig.2B-C**).

**Figure 2:**
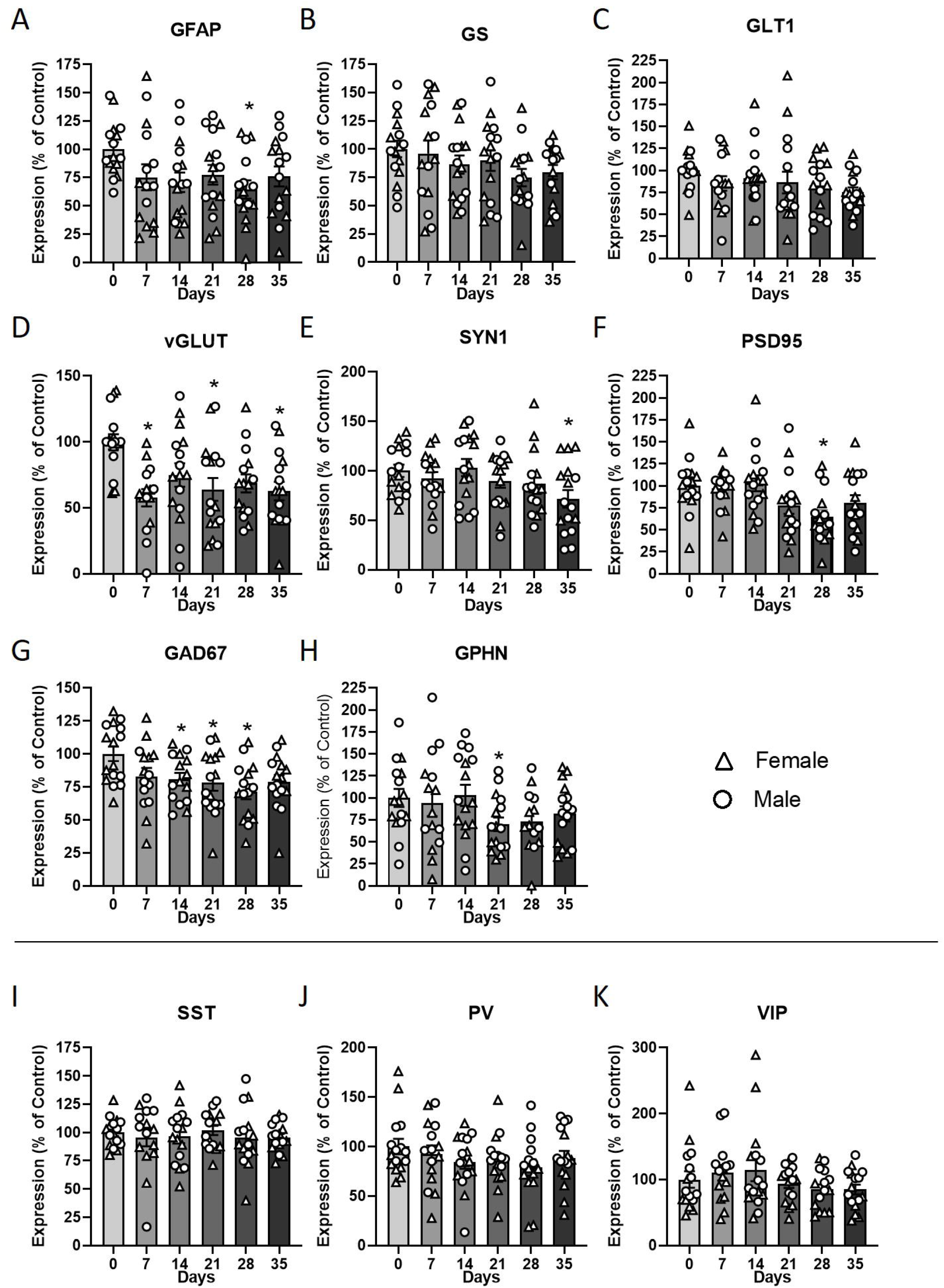
Chronic stress induces gradual emergence of molecular changes in the PFC. Two hours after the last stressor, brains were collected and dissected to collect the prefrontal cortex. Protein and RNA were extracted. Using Western Blot, we quantified the expression levels of GFAP, GS, GLT1, vGLUT, SYN1, PSD95, GAD67 and GPHN (A-H). Using qPCR, we quantified the expression levels of SST, PV and VIP (I-K). *p<0.05, **p<0.01 and ***p<0.001 compared to controls (CRS0).

ANOVA of vGLUT1 protein levels revealed a significant main effect of CRS duration (F_(5:82)_=4.1; p=0.002), but no main effect of sex and no CRS duration*sex interaction (**Fig.2D**). Dunnett’s *Post hoc* analysis identified significant decreases in vGLUT1 levels in the CRS7, CRS21, CRS28 and CRS35 groups when compared to controls. Analysis of Syn1 protein levels showed a significant main effect of CRS duration (F_(5:82)_=2.43; p=0.04) and sex (F_(1:82)_=12.33; p<0.001) but no CRS duration*sex interaction. We identified an overall significant decrease in Syn1 protein levels in the CRS35 group when compared to controls (**Fig.2E**). Splitting by sex, Dunnett’s *Post hoc* analysis revealed a significant decrease of Syn1 expression in the CRS35 group in males (**Supplementary Fig.6E**), while the females showed no significant effect between groups. In addition, Syn1 protein levels were significantly higher in males compared to females in the CRS7 and CRS35 groups. Statistical analysis of PSD95 protein levels indicated a significant main effect of CRS duration (F_(5:82)_=3.12; p<0.05) but no main effect of sex or CRS duration*sex interaction (**Fig.2F**). *Post hoc* analysis revealed a significant decrease in PSD95 levels in the CRS28 group when compared to controls. Analysis of GAD67 protein levels showed that one sample from the CRS14 group was considered a significant outlier compared to the rest of the group and was removed from statistical analyses. ANOVA of GAD67 protein levels showed a significant effect of CRS duration (ANOVA: F_(5:81)_=2.9; p=0.017), and no effect of sex (p=0.28) or CRS duration*sex interaction (**Fig.2G**). The main effect of CRS duration was due to significantly reduced GAD67 levels in the CRS14, CRS21 and CRS28 groups. Analysis of GAD67 protein levels showed a significant effect of sex (p=0.01), but no effect of CRS duration or interaction between factors (**Supplementary Fig.6G**). Analyses of GPHN expression levels showed that one sample from the CRS28 group was a significant outlier and was removed from downstream analyses. ANOVA showed no effect of CRS duration (F_(5:81)_=1.7; p=0.127), and an effect of sex (p=0.02) and no CRS Duration*sex interaction (Fig.2H). The main effect of sex was due to an overall reduced expression of GPHN in females compared to males (**Supplemetary Fig.6H**).

Using qCPR, quantification of expression of SST, PV and VIP mRNA was measured. Statistical analysis of SST mRNA levels indicated no significant effect of CRS duration, sex or CRS duration*sex interaction (F_(5,82)_<1.9; p>0.01; **Fig.2I**). ANOVA of PV mRNA expression levels identified a significant effect of sex (F_(1,82)_=8.6, p=0.004) but not effect of CRS duration or interaction (F_(5,82)_<1.7; p>0.1; **Fig.2J**). The main effect of sex was explained by significant decrease of PV in females in the CRS28 and CRS35 groups, compared to males (**Supplementary Fig.6J**). Finally, ANOVA of VIP mRNA expression levels showed no significant effect of CRS duration (F_(5,82)_=1.4, p=0.2; **Fig.2K**) but a significant effect of sex (F_(1,82)_=3.8, p=0.05) and an CRS duration *sex interaction (F_(5,82)_=2.9, p=0.016). *Post hoc* analyses revealed a significant higher expression level in male in the CRS7 and CRS28 groups compared to females.

### Cellular markers correlate with behavioral outcomes

Relationship between marker expression and behavioral outcomes was assessed using correlation analysis (**Fig. 3** and **Supplementary Table 3**).

**Figure 3:**
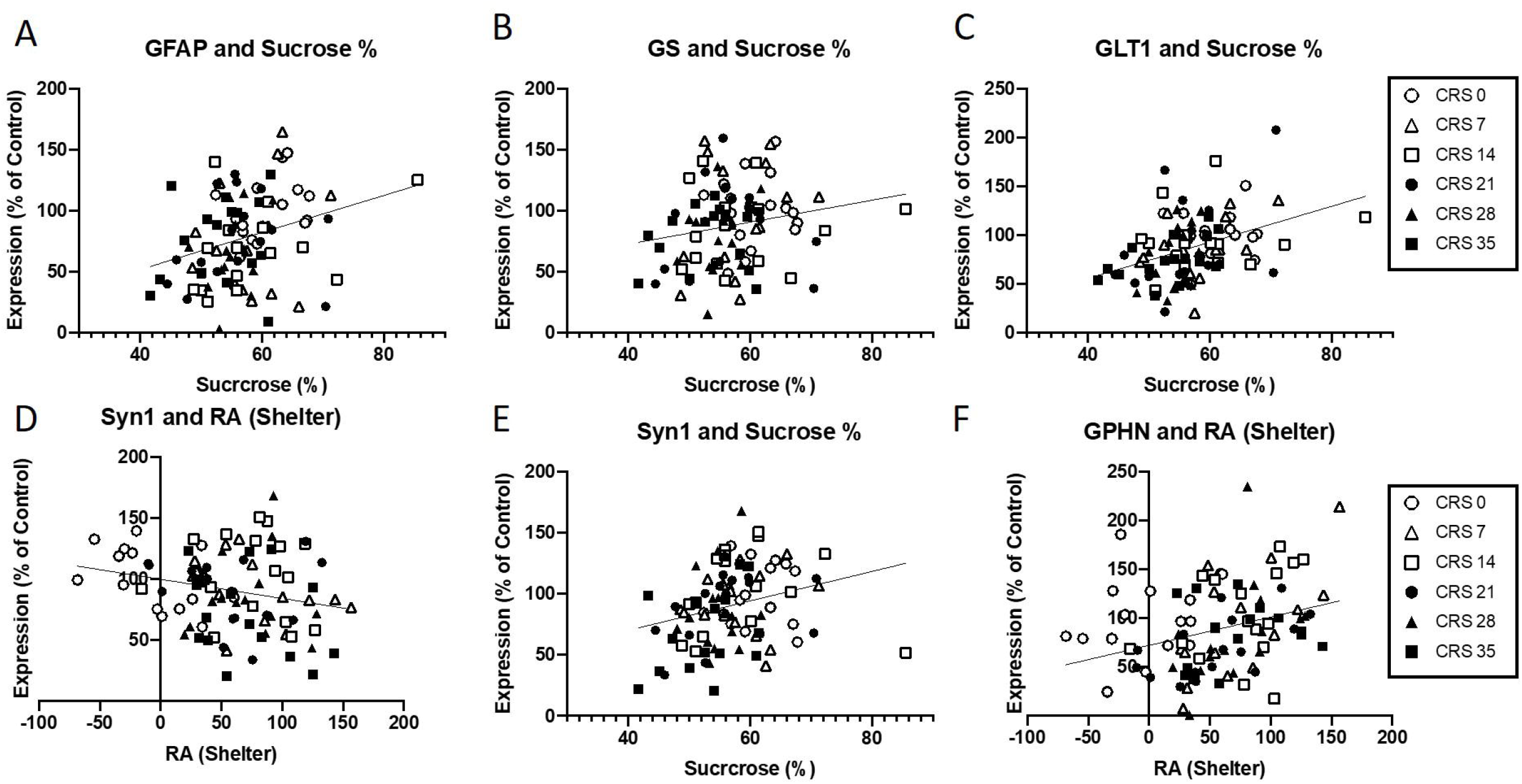
Correlation between molecular changes and behavioral outcomes. Pearson’s regression analyses between expression levels of molecular markers and behavioral outcomes were performed. Sucrose consumption correlated positively with expression levels of GFAP (A), GS (B) and GLT1 (C). Expression levels of SYN1 also correlated with the RA shelter (D) and the sucrose consumption (E). Expression levels of GPHN also correlated with the RA shelter (F).

Overall astroglial marker expression positively correlated with anhedonia-like behaviors. Indeed, we found significant correlations between sucrose consumption and GFAP (R=0.4, p<0.001; **Fig. 3A**) or GLT1 (**Fig. 3B**; R=0.3, p=0.003) expression levels. Pearson’s regression analyses also revealed trends between GS expression levels and sucrose consumption (R=0.19, p=0.07) and RA SZ (R=-0.19, p=0.07). Interestingly, splitting the dataset by sex showed different results. In female mice, only GLT1 expression level correlated with sucrose consumption (R=0.4, p=0.006). In male mice, all glial markers correlated with sucrose consumption. Exclusively in male mice, GFAP and GLT1 expression levels correlated with weight gain, and RA-SZ (p<0.04; **Supplementary Table 3; Supplementary Figure 7**).

Analyses of synaptic markers showed that Syn1 expression levels correlated with RA-SZ (R=-0.23, p=0.02) and sucrose consumption (R=0.26, p=0.008; **Fig.3D-E**). Using Spearman’s regression, we found that vGLUT1 expression levels correlated with the coat state score (Rho=-0.33, p=0.001). When spitting the dataset by sex, we found similar correlation in male mice and no correlation in female mice. More precisely, in male mice, the correlations identified between Syn1 expression levels and, RA-SZ and sucrose consumption, as well as correlation between vGLUT1 expression levels and coat state score were all maintained. In addition, PSD95 expression levels correlated with sucrose consumption (r=0.32, p=0.02), only in male mice.

Focusing on GABA markers, Spearman’s regression analyses on GAD67 expression levels identified negative correlation with coat state (Rho=-0.4, p<0.001). Overall GPHN expression levels were also significantly correlated with RA-SZ (R=0.3, p=0.002, **Fig.3 F**), and coat state score (Rho=-0.22, p=0.03). Overall VIP expression level correlated with weight gain (R=0.24, p=0.018). As previously observed, correlation between markers and behavioral outcomes were more maintained in males than in females. Indeed, only correlation between the coat state scores and GAD67 (Rho=-0.4, p=0.007) and GPHN (Rho=-0.5, p=0.017) expression levels remained in female mice after splitting by sex. In male mice, GAD67 expression levels correlated with coat state score (Rho=-0.38, p=0.01), and SST expression levels correlated with weight gain (R=0.35, p=0.01) and RA-SZ (r=-0.33, p=0.026).

Finally, possible correlations between all markers and z-scores were investigated (**Fig.4**). Z-score correlated with GLT1 (R=-0.3, p=0.004), vGLUT1 (R=-0.27, p=0.008), Syn1 (R=-0.32, p=0.001), GAD67 (R=-0.3, p=0.002) and was trending towards significance with PSD95 (R=-0.18, p=0.08). After splitting by sex, different profiles emerged for males and females (**Supplementary Figure 7–9**). In males, significant correlation were found with GLT1 (R=-0.4, p=0.0004), GFAP (R=-0.29, p=0.04), Syn1 (R=-0.36, p=0.001), GAD67 (R=-0.3, p=0.01) and was trending towards significance with vGLUT1 (R=-0.26, p=0.07) and PSD95 (R=-0.18, p=0.06). In females, only vGLUT1 was significantly correlated with the z-score (R=-0.4, p=0.005), while GPHN was trending towards significance (R=-0.25, p=0.08).

**Figure 4:**
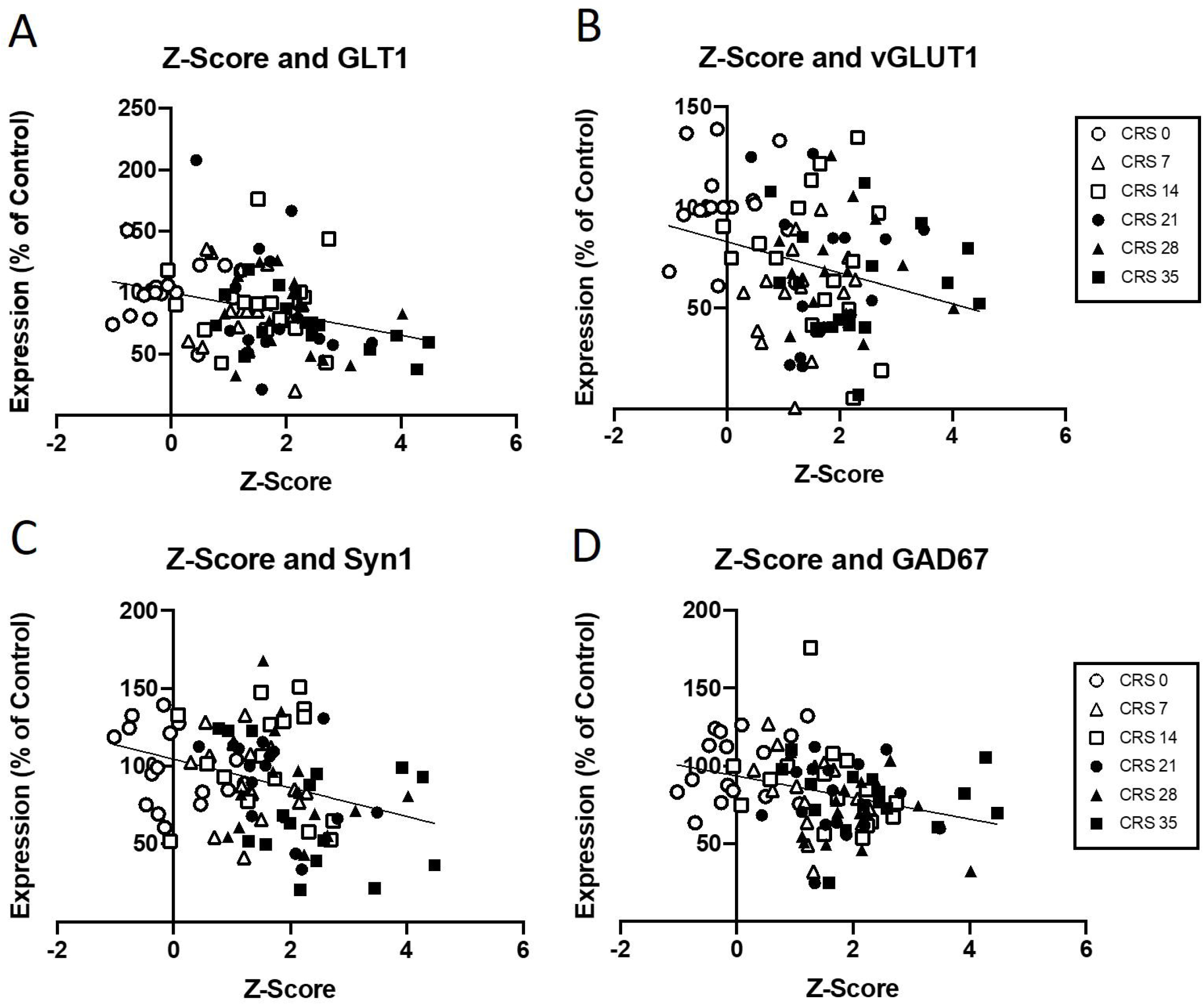
Correlation between molecular changes and Z-Scores. Pearson’s regression analyses revealed that Z-score negatively correlated with GLT1 (A), vGLUT1 (B), Syn1 (C), GAD67 (D).

### Co-expression and principal component analysis

Markers expression levels were also correlated with each other (**Supplementary Fig.10** and **Supplementary Table 4**) and in sex-dependent manner (all described in more details in the **Supplementary Results**). PCA and network co-expression analyses were used to interpret the broad patterns of marker expression in the data across durations of stress. First, we performed a simple pairwise correlation analysis, which showed that most markers were positively correlated with each other, with the glial markers GLT1 and GFAP, as well as GLT1 and GS, and GFAP and GS being particularly strongly correlated with each other (Figure 5 A and Supplementary Table 4). GS and GPHN were the only markers that were significantly negatively correlated with each other (Figure 5A).

**Figure 5:**
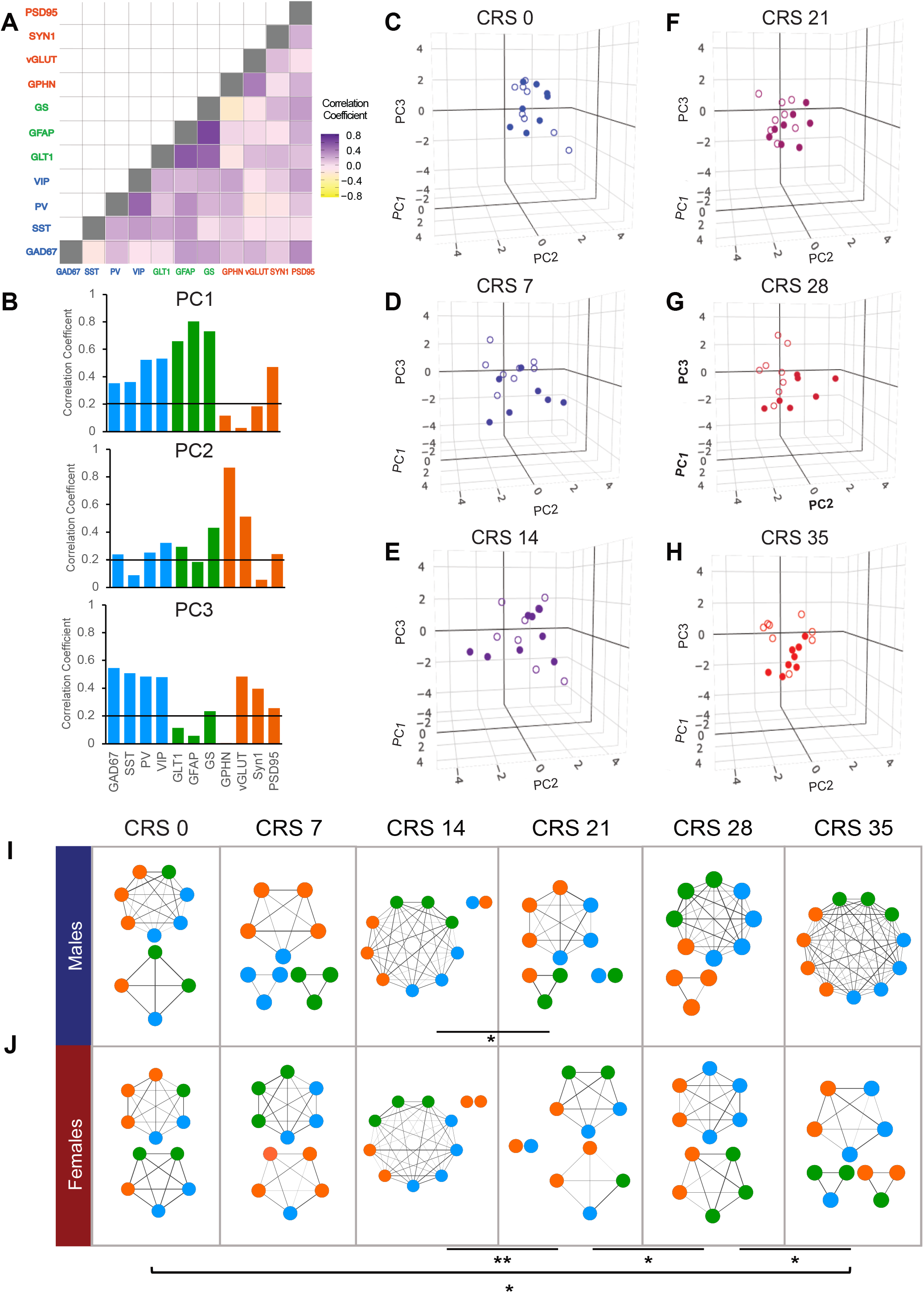
Marker expression as analyzed through pairwise correlations, principal component analysis, and network co-expression analysis. **A**) Marker x marker pairwise Pearson correlation r values combining both sexes and all stress groups. **B**) Correlation of marker loadings with the first three principal components. The horizontal black line marks the critical r-value to reach statistical significance for 55 unique pairwise comparisons (α =0.05). **C-H**) Individual mice plotted in 3-D along the first three principal components. The axes have been oriented such that, progressing through the plots from CRS 0 – CRS 35, differential migration between the sex groups is evident when tracking points from the top right towards the bottom left. **I-J**) Co-expression analysis network diagrams for the male and female mice. *Asterisks represent statistical significance after Benjamini-Hochberg FDR correction, where * denotes q < 0.05; ** denotes q < 0.01. In 5A, 5B, 5I, and 5J, blue denotes GABAergic markers, green denotes astroglial markers, and orange denotes synaptic markers. In 5C-H, filled circles denote males and open circles denote females*.

Correlation matrices between the molecular markers were used to perform a principal component analysis (**Fig.5**). Together, the first 5 PCs accounted for about 75% of the total variance in the data with the first three PCs explaining 24.5%, 14.6%, and 14.1% respectively. Correlation of the loadings of each marker with PC1, PC2, and PC3 elucidated the weighting of the GABAergic, astroglial, and synaptic markers on each of these components (**Fig.5B**). PC1 primarily loaded the astroglial and GABAergic markers, while PC2 loaded the majority of the synaptic markers, and PC3 loaded a portion of the GABAergic and synaptic markers. Due to the relatively equal weighting of PC2 and PC3 on the variance explained, we plotted three-dimensional scatterplots in order to visualize the data for each group (**Fig.5C-H**). Notably, there appears to be a sex-specific progressive temporal effect of stress that results in a left and downward shift, which seems to distinguish males and females after 28 and 35 days of CRS (**Fig.5G,H**).

None of the PC correlated with z-score. We also analysed potential relationship between each PC and each behavior and found that only PC1 significantly positively correlated with sucrose consumption (r=0.34, p < 0.001).

### Glial, GABAergic and synaptic networks alterations by chronic stress are sex-dependent

Behavioral and molecular analyses suggested potential sex-dependent changes due to chronic stress exposure. Indeed, it seems that while female showed more significant molecular changes, correlation analyses revealed that males display more significant correlation between behavioral and molecular outcomes. To further investigate the effect of stress duration on the expression of the GABAergic, astroglial, and synaptic markers in a sex-dependent manner, network co-expression analysis was performed. We found a notably different reorganization of markers in response to stress in males and females (**Fig.5I-J**). While both sexes had significant changes in the makeup of networks between stress durations, only the females had a significantly different network reorganization at CRS 35 when compared to the control group (p = 0.0078, q = 0.014), where the network seems to become more disorganized after CRS. The females also show significant network changes between each week from CRS 14 onward; more specifically, we found group differences in module composition between CRS 14 and CRS 21 (p = 0.021), CRS 21 and CRS 28 (p = 0.0054, q = 0.0096), and CRS 28 and CRS 35 (p = 0.01, q = 0.018) (Figure J). The males appear to undergo a network reorganization which results in a highly organized single network of all the markers at CRS 35 (**Fig.5I**), however this effect was not significant (p =0.099). Notably, the males do show an early reorganization in response to stress between CRS 14 and CRS 21 (p = 0.013) (**Fig.5I**). Altogether, this suggests that while behavioral and molecular changes observed are fairly the same in males and females, underlying network connectivity and contribution to the pathology is sex-dependent.

## Discussion

Chronic stress exposure is linked to various brain physiology changes, such as changes in excitation, inhibition and support of the brain cells. While the behavioral and brain changes are well characterized, the trajectory of such changes remain unclear. Here, we investigated the impact of various duration of chronic stress exposure in a mouse model known to ultimately induce behavioral and molecular/cellular changes relevant to human MDD. We demonstrated that anxiety-like behaviors are the first behavioral response to emerge during chronic stress exposure, followed by anhedonia-like behaviors with longer exposure. At the molecular level, we demonstrated that chronic stress induces a progressive reduction in markers of function and structure of astrocytes, GABAergic interneurons and excitatory synapses. Using correlation analyses coupled with principle component analyses and network analyses, we showed that the impact of CRS is dependent on its duration, and that, while showing fairly similar behavioral and molecular changes, male and female mice have distinct responses to CRS considering network organization mediating the impact of CRS.

In this study, we confirmed that chronic stress exposure reduces behavioral changes, as early as 1 week after initiation of the chronic stress paradigm. We show that anxiety-like behavior are present from week 1 and onward, using the Phenotyper tests developed by our group Prevot et al.[44] The results obtained in the present study are consistent with previous findings reported using the Phenotyper test, and further confirm the validity of this test to detect anxiety-like behaviors in a repeatable fashion. Our results are also consistent with other findings showing that chronic restraint stress induces exacerbated anxiety-like behaviors in rodents [54, 55]. Guedri et al. [55] previously showed that CRS in rats, 3hrs per day for 14 days induces anxiety-like behaviors observed in the open field and the elevated plus maze. Chiba et al. [54] also showed that CRS in rats, 6hrs per day for 28 days also induces anxiety-like behaviors. Our data in mice, using the Phenotyper test to assess anxiety-like behaviors repeatedly confirmed these results, in mice and in a repeatable fashion from 7 to 35 days of CRS. While anxiety-like behaviors were observed throughout the duration of the CRS paradigm, anhedonia-like behaviors emerged more gradually, and were detected only after 35 days of chronic stress, which is in accordance with other studies showing that chronic stress exposure in mice or rats induces anhedonia-like behaviors [56]. Other studies suggested that chronic mild stress, but not chronic restraint stress can induce anhedonia-like behaviors in mice [57]. However, these studies uses paradigm that last less than 28 days, while ours lasted 35 days. This shows how critical the duration of the chronic stress is when studying the impact of chronic stress on behavioral outcomes, and suggests that shorter chronic stress duration may not be able to detect anhedonia-like behaviors, compared to longer paradigms. Considered altogether using a z-score approach, we see that CRS induces behavioral changes throughout the weeks of chronic stress exposure, but it appears critical to still consider the individual behavioral outcomes, as they suggest various trajectory for anxiety-like behaviors and anhedonia-like behaviors. Indeed, as mentioned before, the shorter durations of chronic stress exposure induced anxiety-like behaviors (Week 1-2-3), while the longer duration induced anxiety-like behaviors as well as anhedonia-like behaviors (Week 5). One could think that there is a biphasic mechanism in place that could represent a pseudo-adaptive response at first, still marked by anxiety-like behaviors, that transforms into a more pathological response which combines anxiety-like behaviors and anhedonia-like behaviors. This is in part confirmed by our network analyses, where male and female mice seem to develop a similar response to CRS in the early stage of exposure (up to 2 weeks), while a more complex network reorganization is observed in the late stages (4-5 weeks).

This observation is also supported by the reduced effects of chronic stress on molecular markers in the early stages of chronic stress exposure. Indeed, after 7 days of CRS, only expression levels of vGLUT1 in the PFC were significantly reduced. The fact that vGLUT1 is the first and only marker to be significantly reduced after 1 week of CRS is consistent with the results from previous studies showing that reduced levels of vGLUT1 in the PFC is responsible for increased susceptibility to chronic stress [58] and facilitate the emergence of anxio-depressive-like behaviors and neuroendocrine responses in rodents [59]. With longer CRS exposure (14-21 days), we observed that the expression levels of GAD67 and GPHN were significantly reduced. These results are again consistent with previous studies similar reduction in other models of chronic stress paradigms [60–62], and in depressed patients [63]. Finally, in later stages of CRS exposure (28-35 days), we saw that expression levels of vGLUT1 were still lowered, and we observed reduced expression of GFAP followed by reduced expression of SYN1 and PSD95. Reduced levels of GFAP expression after chronic stress exposure is well characterized [64–67], and our results are consistent with what has been shown in the past. Interestingly, one could see that the diminution in expression level of GFAP decreases early on during CRS exposure, but that the inter-individual variability is high. Looking at sex differences, it is also noticeable that female have overall reduces expression levels of GFAP throughout the CRS exposure period, highlighting a first sex difference. The other glial markers investigated did not reach significance for changes induced by CRS. Expression levels of GS were not significantly affect by CRS, consistent with previous findings [67] using chronic unpredictable stress in rats. Changes in expression levels of GLT1 did not reach significant either, but previous findings showed decreased expression levels in the hippocampus after chronic stress [68, 69]. Here, it is possible that the inter-individual variability that we can observe is the reason why we do not find significant decrease in GLT1 expression levels after CRS, and this is observed in both males and females. However, at 35 days, inter-individual variability seems to be reduced and it could be plausible that GLT1 expression levels would have reached significant if CRS was applied for one more week.

Synaptic markers SYN1 and PSD95 showed significant decrease after 35 and 28 days of CRS, respectively. Previous studies found similar results with reduced expression levels of SYN1 in the PFC or HPC after chronic stress, as well as reduced expression levels of PSD95 even after chronic corticosterone administration [70]. These changes are region dependent, as other studies showed that unpredictable chronic mild stress induces an increase in PSD95 expression level in the amygdala [71], consistent with increased emotionality and amygdala-centered activity in chronic stress state and MDD [71].

Analyses of the expression levels of the GABAergic markers showed a significant reduction in GAD67 and GPHN, consistent with previous studies showing similar reduction in other models of chronic stress paradigms [60–62], and in depressed patients [63]. Here, we did not find significant reduction in SST, PV or VIP mRNA levels, which is inconsistent with previous findings. Indeed, SST levels are known to be reduced in MDD patients [32, 72], and in animals subjected to chronic stress [73]. Regarding expression of PV, previous studies reported a decrease in expression in the dorsal HPC after chronic stress [74], but an increased expression in the PFC, associated with increased function exacerbating anxiety-like behaviors in mice [75]. In the study by Page et al. [75], the increased activity of the PV cells in the PFC was also more important in females than males, using chemogenetic approach increasing specifically the activity of the PV cells. Interestingly, while in opposition, we showed here that there is a significant sex-difference in PV expression levels in the PFC of mice subjected to 28 and 35 days of CRS. Indeed, female mice showed reduced level of PV expression, while male expression levels were unchanged. One could think that the impact of chronic stress on PV cells is sex-dependent and also depends on the paradigm used.

Using regression approaches, we found that glial markers (GFAP, GS and GLT1) correlated positively with sucrose consumption, suggesting that animals with reduced expression of glial markers would exhibit anhedonia-like behaviors. Previous studies showed similar relationship between glial markers and anhedonia [76, 77]. The synaptic marker SYN1 strongly correlated with anxiety-like readouts, as well as with sucrose consumption, suggesting that reduced levels of SYN1 expression are linked to increased anxiety-like behaviors and anhedonia, consistent with previous findings [78]. Again, such correlation analyses were found stronger in males than in females in the present studies. Finally, GABAergic markers correlated with anxiety-like behaviors exclusively, and were again stronger correlated in males than females. Such changes in the GABAergic system and their link to anxiety-like behaviors are consistent with previous finding, where reduced GPHN levels in the hippocampus are proposed to contribute to anxiety-like phenotype [79, 80]. Similarly, loss of GAD67 function in particular in SST cells is proposed to contribute to anxiety-like phenotype[81] and reduced levels of GAD are reported in human MDD with comorbid anxiety disorders[82, 83].

Interestingly, while clinical and preclinical studies have shown higher sensitive to chronic stress in females compared to males, here we found stronger correlation between glial markers and behavioral outcomes in males, compared to females. This seems to be driven by the greater amplitude of the behavioral outcomes observed in males, but could also be the result of a more complex network reorganization in females. This is supported by our results. Indeed, when we observe the network makeup in males and females, we see that for the 2 first weeks of CRS exposure leads to similar early network re-organization in both sexes, ultimately increasing connectivity between the nodes. This suggests a common response to early stress exposure. Then, we see that network organization at CRS21 is different to the previous time point in both sexes, suggesting a major change after the 2-3 weeks milestone. At the latest time points, we see a clear dissociation in network organization where males tend to develop a single network, highly connecting the different nodes, while females seem to have 2 to 3 sub-networks independent from each other. This suggest that the underlying network connectivity and contribution of each compartment to the pathology is sex-dependent, complicating our understanding of the underlying mechanisms underlying symptoms expression and reducing treatment intervention, consistent with what is observed in human MDD.

This study also presents some limitations. To start with, we only focused on the PFC, while other brain regions are impacted by stress or chronic stress like the hippocampus or the amygdala to only cite a few. It could be of interest to perform the same study in the hippocampus and the amygdala in order to investigate the gradual impact of CRS in these brain regions and complement our network analyses in a more integrative fashion. Also, we used chronic restraint stress as a model to induce behavioral and molecular changes in mice while other studies use chronic unpredictable mild stress. This choice was driven by the proof of efficacy of the CRS model and by the fact that with such a high number of animals in our study. CRS was a better choice than UCMS for such a time course study, as UCMS is known to rely on variability between stressors to induce behavioral and molecular changes, themselves subjected to variability between studies, therefore counter indicated in our case. Another limitation is that we limited our investigation of the 3 compartments (astroglia, GABA cells, and synapse) to key markers. Indeed, we could not perform Western Blot and qPCR on all specific astroglial, GABAergic and synaptic markers that exist. A follow up could investigate additional markers, or could perform RNA-sequencing to determine gene-expression changes caused by CRS duration. Finally, our analyses were performed on samples from bulk PFC. We did not use a single-cell approach to characterize the protein and RNA expression of astroglial cells, GABA cells and glutamatergic neurons individually. With the advances regarding RNA scope, it would now be feasible to perform qPCR or RNA-seq, on single cells collected on a laser-captured microdissection microscope to refine our analyses of the compartment-specific changes.

To conclude, we show that chronic stress induces behavioral and molecular changes in a time-dependent, compartment-dependent, and sex-dependent manner. We highlight that shorter duration of CRS exposure may be models of acute to “adaptive” stress response, while longer chronic stress exposure may induce more “pathologically-relevant” behavioral and molecular changes, aligned with what can be observed in human MDD. We also highlight that while apparently similar in symptom expression and cellular changes, females present a distinct underlying network organization that may explain what is observed in human MDD, i.e. more severe symptoms and reduced efficacy of treatment.

## --- Supplementary Information ---

### Supplementary Materials and Methods

#### Animals

In this study, we used 8 week-old C57BL/6 mice (n=96) from Jackson Laboratories (Stock No: 000664; Bar Harbor, Maine, USA). Prior to experimentation, animals were habituated to the facility for 1 week with a 12h light/dark cycle and *ad libitum* access to food and water. The animals were assigned either the control group or a CRS group in which they were subjected to 7, 14, 21, 28 or 35 days of CRS (n=16 animals/group, 50% females). The control group was handled daily for 3 days and given nesting material (Ghosal et al., 2015). Control mice were housed in a separate room than the CRS groups to avoid any disruption. All experiments were conducted in line with guidelines provided by the Canadian Animal Care Committee and protocol was approved by the animal care committee of the *Centre for Addiction and Mental Health*.

**Coat State and Body Weight** were measured weekly. Coat state was assessed following the method described by Yalcin et al.[45]. Seven body parts were assessed for overall appearance of the coat (head, neck, dorsal coat, ventral coat, tail, forepaws, and hind paws). Each body part was given a score of 0 for a well-groomed coat, 0.5 as an in-between, or 1 for a deteriorated coat. The sum of the scores for all the body parts was considered to be the overall coat state score of the animal. Weight gain was measured using the weight of each mouse on Week 0 as the reference.

#### Sucrose Consumption

This test was also performed on a weekly basis. Mice underwent a 48hr habituation period to a 1% sucrose solution (only for the first exposure – subsequent habituation only lasted 24hrs). Then the mice were deprived of any liquids for a 16hr overnight period (^~^6pm to 10am). On the subsequent morning, the sucrose solution was returned to the mice for a 1hr period after which consumption was recorded. This same protocol is repeated as a measure water consumption to be used for comparison. Each week the ratio of consumption was analyzed as ratio of consumption which is calculated as: 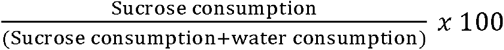.

##### PhenoTyper

The PhenoTyper test was used on a weekly basis to assess anxiety-like behavior as per the protocol described in Prevot et al., 2019. During this test, animals were placed in the PhenoTyper^®^ (Noldus) boxes (30cm^2^ arena) that are set up similar to a home cage with a food dispenser, water bottle and shelter. An infrared camera, mounted in the ceiling of the box, recorded the animal’s activity. Ethovision^®^ software was used to track the time spent in the arena and in specific zones: the food zone and shelter zone. Animals were placed in the PhenoTyper^®^ (Noldus) at the beginning of the evening (6pm), and were left in the box overnight, until 9am the next morning. Baseline activity was recorded from 7pm to 11pm. At 11pm, a light challenge was applied over the food zone, for 1hr. After the one-hour period of aversive stimulation, the light turns off. Overall activity is recorded during baseline, during the light challenge and after the light challenge. The information recorded using Ethovision^®^ software can then be used to calculate the Residual Avoidance (RA) score, as outlined below.

##### Residual Avoidance Calculation

We calculated the Residual Avoidance (RA) of time spent in the food zone (FZ) and shelter zone (SZ) during the PhenoTyper test as a proxy for anxiety-like/depressive-like behavior. This measure had been previously validated in our lab and peer reviewed in the paper Prevot et al., 2019. This metric represents how animals react after the light challenge as compared to controls. The RA metric can be expressed as:

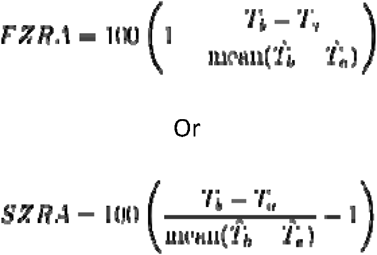

Where *T_b_* is the time spent in the zone (either FZ or SZ) from 12am-5am, *T_a_* is the time spent in the zone from 11pm-12am and ^ over top signifies the control group. If the mouse avoids the illuminated zone by either avoiding the food zone or spending the time hidden in the shelter, then RA>0. Average of the RA from the control animals is always 0. This RA calculation informs us about the reaction after the mice after light challenge as compared to controls. RA scores were calculated in a sex-dependent manner, with the average of the RA from the male and female control groups, both being equal to 0.

#### Sample collection and preparation

Twenty-four hours after the last stressor, brains were collected, PFC was dissected and frozen using dry ice. Using a Qiagen Allprep RNA/protein Kit (#80404), RNA and proteins were extracted. RNA was converted into cDNA using a SuperScript VILO cDNA Synthesis Kit (ThermoFisher, Massachusetts, USA, Cat#: 11754050). Proteins extracted from each PFC sample was quantified using the Pierce BCA (Bicinchoninic Acid) Protein Assay Kit (Thermofisher, Massachusetts, USA, Cat #: 23250)

#### Western Blot

20 μg of protein per sample was loaded into BioRad Criterion TGX Stain-Free Precast gel (4-20%). Electrophoresis was then run at 70V for 1 hour. Total protein was imaged and quantified using a molecular imager (ChemiDoc XRS, BioRad) with ImageLab™ software, directly in the gel, using the TGX Stain-Free technology. Proteins were transferred from the gel onto a nitrocellulose membrane which were then incubate with the primary antibodies (GLT1, GFAP, GS, Syn1, vGLUT, PSD95, GAD67, and GPHN) at 4°C overnight. Membranes were then washed and incubated but with the 2° antibody for 1hr at room temperature (see **Supplementary Table 1**). Enhanced chemiluminescence (ECL) substrate was used for detection and then signal was analyzed with Image Lab™ software.

#### cDNA synthesis and qPCR

Using cDNA samples, standard qPCR experiments were performed to amplify cDNA of somatostatin (SST), parvalbumin (PV) and vasopressin (VIP). 1 μL cDNA, SsoAdvanced Universal SYBR Green Supermix (BioRad, California, USA; Stock Number: 64190925), and respective primers are placed into the BioRad CFX96™ Real-Time System C1000 Touch™ Thermal Cycler qPCR Machine (BioRad, California, USA; Stock Number: 1855195). Data were analysed using BioRad CFX Manager 3.1 Software (3.1.1517.0823) All primers used were from Integrated DNA Technologies (IDT; Iowa, USA):

- Actin (F: 5’-CCTAGCACCATGAAGATCAA-3’; R: 5’-GGAAGGTGGACAGTGAGG-3’)
- SST (F: 5’-CAACTCGAACCCAGCAAT-3’; R: 5’-GGTCTGGCTAGGACAACAA-3’)
- GAD1 (F: 5’-CTTCAGGGAGAGGCAGTC-3’; R: GGAGAAGTCGGTCTCTGTG-3’)
- BDNF-CDS (F: 5’-CAGTATTAGCGAGTGGGTCA-3’; R: 5’-CCTTTGGATACCGGGACT-3’)
- GAPDH (F: 5’-AACTCCCACTCTTCCACCT-3’; R: 5’-CACCACCCTGTTGCTGTA-3’)
- SYN1 (F: 5’-ATGCAAACTCCACCCATC-3’; R: 5’-AGGAGGCCAAGTCAGTCA-3’)
- VIP (F: 5’-GACATCTTGCAGAATCCCTTA-3’; R: 5’-CTGCTGTAATCGCTGGTG-3’)
- vGLUT1 (F: 5’-CTATGTCTATGGCAGCTTCG-3’; R: 5’-TCAATGTATTTGCGCTCCT-3’)
- GFAP-Tr1 (F: 5’-CGCATCACCATTCCTGTA-3’; R: 5’-GAGCCTTTTGAGAGGTCTTG-3’)
- GFAP-Tr2 (F: 5’-ATCCGCTCAGTCATCTTACC-3’; R: 5’-CCCTTAGCTTGGAGAGCA-3’)
- PVALB (PrimeTime^®^ qPCR Primers, Mm.PT.58.7596729, Pvalb Exon Location 3-4, 20X [F+R])
- CYCLO (PrimeTime^®^ qPCR Primers, Mm.PT.39a.2.gs, Ppia Exon Location 4-5, 20X [F+R]).

Each sample was run in quadruplicates. Results were normalized to three validated internal controls (actin, GAPDH, and cyclophilin G) and calculated as the geometric mean of threshold cycles. The results are further expressed as a percentage of control group.

#### Network Analysis

Co-expression analysis was used to examine changes in coordinated expression between the markers. All analyses were performed in R [46], version 3.6.0. Analytical code is available upon request. A similar approach to gene co-expression, module generation, and preservation analysis was taken as in a recent study examining co-expression changes in a similarly sized cohort [47]. Briefly, within each group we generated Pearson correlation matrices of all markers, Z-normalized to account for marker scaling differences. Markers were then hierarchically clustered based on degree of correlation, and modules were generated from the resulting dendrogram using the dynamicTreeCutting function from the WGCNA R package [48]. A minimum module size of three was used, as a two member module simply reflected a pairwise correlation. Though we employed particular functions from the WGCNA package, we opted not to use the entire workflow as various steps such as topographical overlap matrix generation and matrix multiplication are not appropriate for our limited set of genes. Networks were visualized in Cytoscape [49]. Co-expression modules were compared across stress groups using permutation testing to assess the degree to which they were preserved over the course of CRS. We examined the difference in module composition in each CRS group versus controls, and between each CRS group. We used permutation testing (n=10,000) to compare cross tabulation-based measures of module preservation (i.e. whether markers remained co-expressed or not) and generate distributions of variability in module composition. These distributions were used to generate module-wise empirical p-values. Significant p-values represented a lesser degree of preservation (i.e. coexpression modules showed a different composition). Fisher’s p-value meta-analysis was used to combine module-wise p-values into a single p-value for each group-to-group comparison after Benjamini-Hochberg FDR correction [50].

### -Supplementary Results-

#### Weight Gain

Repeated measure ANOVA of weight gain assessed weekly (**Supplementary Fig.1A**) showed no significant difference between groups (F_(5;450)_=0.69; p=0.62), but showed a significant effect of weeks (F_(5,450)_=37.81, p<0.0001), characterized by significant increase of weight from week to week. To better visualize this effect, weight gain was split per group (**Supplementary Fig.1B-G**). In all groups, the effect of the weeks was confirmed (p<0.004). In control mice, significant increase in weight gain compared to baseline was found after 3 weeks and onward. Similar profile was found in the CRS7 and CRS35 groups. In the CRS14 and CRS21 groups, weight gain was significantly higher than baseline from week 1 and onward. Finally, in the CRS28 group, weight gain was significantly higher than baseline only in week 4 and week 5. ANOVA of weight gain at week 5, across groups (**Supplementary Fig.1H**) did not reveal significant effect of CRS duration (F_(5,82)_=0.64, p=0.66), sex (F_(1,82)_=0.052, p=0.82) or interaction (F_(5,82)_=0.51, p=0.76).

#### Coat State Score

Repeated measure ANOVA of the coat state scores over the weeks (**Supplementary Fig.2**) showed a significant effect of CRS duration (F_(5,420)_=33.2, p<0.001), a significant effect of sex (F_(1,420)_=13.27, p<0.001), and a significant effect of weeks (F_(5,420)_=89.71, p<0.001). The main effect of CRS duration is characterized by increasing coat state scores with increasing CRS duration. The main effect of sex is due to higher scores in males than in females (data not shown). Finally, the main effect of weeks is due to increasing scores with weeks, mostly driven by animals subjected to CRS. Indeed, in mice subjected to CRS for 1 week only (CRS7), coat state score on week 5 were higher than baseline (p<0.001), while the scores from week 1, 2, 3 and 4 were not different from baseline. This was also observed in the other groups, characterized by increasing score after beginning of CRS.

#### Weekly testing in the Phenotyper Test

Following the design presented in **Figure 1A**, all mice were tested in the Phenotyper test on a weekly basis, whether they were subjected to CRS or not yet. As an example, mice from the CRS7 group were tested in the Phenotyper test every week, but were only subjected to stress the last week of the experiment (week 4 to 5), meaning that from baseline to the “Week 4” time point, they were technically similar to control mice. **Supplementary Figure 3** shows the time spent in the shelter zone of the Phenotyper, in all mice from baseline to Week 4 (Week 5 being presented in **Figure 1D**). Repeated measure ANOVA of time spent in the shelter at baseline, showed no main effect of CRS duration between groups (artificially defined at this stage: F_(5,504)_=0.1, p=0.99). A main effect of time was found (F_(12,504)_=66.92, p<0.001), and is characterized by increased time spent in the shelter when the light is ON from 11pm to 12am, and then increasing from 3am to 7am (as demonstrated in Prevot et al. 2019). After determination of baseline, animals from the CRS35 group started to be subjected to CRS. On Week 1, only the CRS35 group was subjected to CRS, and at this stage, for only 1 week, while the other groups remained as controls (**Supplementary Figure 3B**). Repeated measure ANOVA of time spent in the shelter on Week 1 showed for the first time a main effect of CRS duration (F_(5,504)_=3.6, p=0.008), a significant effect of time (F_(12,504)_=86.06, p<0.001) and an interaction between factors (F_(60,504)_=1.54, p=0.007). *Post hoc* analyses mainly showed that animals from the CRS35 group spent significantly more time in the shelter compared to control at 2am. On Week 2, animals from the CRS35 group achieved 2 weeks of CRS, while animals from the CRS28 group achieved their first week of CRS. Similarly, repeated measure ANOVA of time spent in the shelter showed a significant effect of CRS duration (F_(5,504)_=2.45, p=0.049), a significant effect of time (F_(12,504)_=92.33, p<0.001) and an interaction between factors (F_(60,504)_=2.04, p=0.007). *Post hoc* analyses on the time spent in the shelter at each time point showed that the CRS28 group spent significantly more time in the shelter compared to control (p=0.009). Similarly, on Week 3, mice from the CRS35, CRS28 and CRS21 groups had been subjected to 3, 2 and 1 week of CRS, respectively. Repeated measure ANOVA of time spent in the shelter showed a significant effect of CRS duration (F_(5,504)_=9.05, p<0.001), a significant effect of time (F_(12,504)_=66.05, p<0.001) and an interaction between factors (F_(60,504)_=3.17, p=0.007). *Post hoc* analyses revealed significant increase in time spent in the shelter, compared to control mice, at 2am and 3 am for mice of the CRS35, CRS28 and CRS21 groups (p<0.05). Finally, on Week 4, mice from the CRS35, CRS28, CRS21 and CRS14 groups were subjected to 4, 3, 2 and 1 week of CRS respectively. Repeated measure ANOVA of time spent in the shelter showed a significant effect of CRS duration (F_(5,504)_=4.16, p=0.0037), a significant effect of time (F_(12,504)_=87.72, p<0.001) and an interaction between factors (F_(60,504)_=2.53, p=0.007). *Post hoc* analyses revealed significant increase in time spent in the shelter, compared to control mice, at 2am for mice of the CRS35, CRS28, CRS21 and CRS14 groups (p<0.05). Also, there was a significant increase in time spent in the shelter, compared to control mice, at 3am for mice of the CRS21 and CRS14 groups (p<0.05). These trajectory analysis of the impact of CRS on a weekly basis illustrate how the Phenotyper test captures the impact, and further demonstrates its validity as shown in Prevot et al. 2019.

#### *Correlation analyses between Marker Expression Levels and Behavioral Outcomes* (Supplementary Table 3)

Pearson’s regression analysis showed no significant link overall between GFAP protein levels and weight gain, SZ or FZ residual avoidance. However, similar analyses performed separately in males and females showed that GFAP protein levels and weight gain or SZ residual avoidance negatively correlated in males but not in the females (**Supplementary Table 3**). In addition, we found a significant positive correlation between GFAP protein levels and sucrose preference (**Fig.3A**). Although significant when including both sexes, correlation was significant only in males (R=0.5, p<0.001) and not in females. Spearman’s regression analysis showed no significant correlation between GFAP protein levels and coat state score, even when splitting by sex. Pearson’s regression analysis between GS protein expression levels and SZ RA or sucrose consumption showed a trend level (**Fig.3B**). When split by sex, regression between GS expression level and SZ RA or sucrose consumption were found significant in males, but not in females. Looking at correlation between GLT1 expression levels and sucrose consumption, Pearson’s regression analysis found a positive correlation (**Fig.3C**), which was confirmed separately in males and in females. Focusing on males, Pearson’s regression analyses found significant correlations between GLT1 expression level and weight gain, SZ RA, and a trend with FZ RA. Spearman’s regression analysis also identified a trend towards significance between GLT1 expression level and coat state, in male mice only.

Pearson’s regression analyses were performed between Syn1 expression levels and behavioral outcomes, identifying significant correlations with FZ RA, SZ RA and sucrose consumption (**Fig.3D-F**). When splitting by sex, all three correlations were only conserved in males. In addition, Spearman’s regression analyses performed between Syn1 expression level and coat state scores found an overall trend towards significance, which did not survive splitting by sex. Pearson’s correlation analyses between vGLUT1 expression level and behavioral outcomes did not find any significant correlations, even when splitting by sex. However, Spearman’s correlation analyses between vGLUT1 coat state score was significant, which is maintained even after splitting the dataset by sex (trend level). Pearson’s regression analyses performed between PSD95 expression and behavioral outcome did not identify significant correlations. Splitting by sex, PSD95 expression levels in male mice correlated positively with sucrose consumption.

Spearman’s regression analyses on GAD67 expression levels identified negative correlation with coat state, which remained significant after splitting the dataset by sex. In males, Pearson’s regression analysis found a trend between GAD67 expression levels and sucrose consumption. Looking at GPHN expression levels, Pearson’s regression analyses found significant correlation with FZ RA, and SZ RA (**Fig.3G-H**). Splitting by sex, correlation with SZ RA were trending in males and females.

Spearman’s regression analyses between GPHN expression level and coat state scores found a significant correlation, which remained significant in females but not in males after splitting by sex. Pearson’s regression analyses on SST expression level only showed a trend towards positive correlation with sucrose consumption, which was maintained in males after splitting by sex. Focusing on male mice, SST expression levels also correlated with weight gain and with SZ RA. PV expression levels did not correlate with any behavioral outcome, even after splitting by sex. Pearson’s regression analyses on VIP expression level only showed a significant correlation with weight gain which was maintained as a trend in females after splitting by sex.

Finally, possible correlations between all markers and z-scores were investigated (**Supplementary Figure 7**). Z-score correlated with GLT1, vGLUT1, Syn1, GAD67 and was trending towards significance with PSD95. After splitting by sex, different profiles emerged for males and females. In males, significant correlation were found with GLT1, GFAP, Syn1, GAD67 and was trending towards significance with vGLUT1 and PSD95. In females, only vGLUT1 was significantly correlated with the z-score, while GPHN was trending towards significance.

#### Correlation Analyses Between Marker Expression Levels

Pearson’s regression analyses using all 11 markers found strong correlations between the three astrocytic markers (GFAP, GS, GLT1) (**Supplementary Table 4 and Supplementary Fig.10**). The structural astroglial marker GFAP strongly correlated with the two functional astroglial markers GS and GLT1, **Supplementary Fig.10A and B**, respectively. GS and GLT1 expression levels also correlated strongly with each other (**Supplementary Fig.10C**). GFAP expression levels also correlated significantly with GAD 67, SST and PV expression levels. Interestingly, none of the astroglial markers correlated with the pre-synpatic markers (Syn1 and vGLUT1). However, GS correlated with the post synaptic markers PSD95 and GPHN. GS also correlated with GAD67, and was trending with PV and VIP. GLT1 correlated with SST and was trending with VIP.

Regarding the synaptic markers (PSD95, Syn1, vGLUT1), they did not significantly correlate with each other, but there was a trend toward a positive correlation between PSD95 and Syn1 levels (**Supplementary Fig.10D**). vGLUT correlated with GPHN (**Supplementary Fig.10E**) and was trending with GAD67. PSD95 correlated with GAD67 (**Supplementary Fig.10F**) and VIP and was trending with GPHN.

Finally, GABAergic markers significantly correlated with each other. VIP expression levels correlated with GPHN (**Supplementary Fig.10G**), SST (**Supplementary Fig.10H**) and PV. PV was correlated with SST (**Supplementary Fig.10I**), and was trending with GPHN.

Splitting the dataset by sex, we can see that correlations between markers is sex-dependent (**Supplementary Table 4**). Interestingly, correlations between all glial markers were maintained, even after splitting the dataset by sex. While no significant correlations were found between GFAP and synaptic markers, splitting by sex showed that GFAP correlated negatively with vGLUT, positively with PSD95 and was trending toward positive correlation with Syn1 in male mice. However, in female, only vGLUT correlated with GFAP, but positively, i.e. in the opposite direction than male mice. Also, GFAP expression in male did not correlate with GABA markers. Only trends were observed with GAD67 and GPHN. In females, GFAP expression correlated with GAD67 and SST. In male mice, other glial markers like GS and GLT1 both correlated negatively with GPHN expression, but not in female mice. In female mice, GS and GLT1 both correlated positively with VIP expression, and GLT1 also correlated positively with SST expression.

While synaptic markers did not correlated between each other considering the entire data set, splitting by sex showed a sex-dependent effect. Indeed, in male mice, Syn1 expression level correlated positively with PSD95 expression level. In female mice, Syn1 expression level correlated positively with vGLUT1 expression level. In both male and female mice, vGLUT expression levels correlated positively with GPHN expression levels, confirming the results observed in full dataset. In male mice, PSD95 expression levels correlated positively with GAD67 while this correlation was only trending in female mice. In female mice, PSD95 positively correlated with VIP expression level.

Finally, GABAergic markers did not correlate between each other in male mice. However, in female mice, GPHN correlated positively with SST, PV and VIP. In female mice, VIP also strongly correlated positively with SST and PV.

Altogether, analyses of the correlation between markers shows that expression levels between compartments are linked, and suggests that there is a sex-dependent relationship between markers.

### - Supplementary Tables -

**Supplementary Table 1:** List of antibodies for western blot analysis. List of primary (1°) antibodies, blocking solutions, and the concentrations used for the western blot analysis for both 1° and secondary (2°) antibodies.

|  | ***Antibody*** | ***Blocking*** | ***Dilution*** | ***Reference*** |
| --- | --- | --- | --- | --- |
| 1° | Rabbit α-GLT1 (EAAT2) | 5% Milk | 1:2,000 | Santa Cruz Biotechnology sc-15317<br>(Mississauga, Ontario, Canada) |
| | Rabbit α-GFAP | 5% Milk | 1:5,000 | Cell Signaling Technology 12389<br>(Danvers, MA, USA) |
| | Rabbit α-GS | 5% Milk | 1:5,000 | Abcam ab176562<br>(Toronto, Ontario, Canada) |
| | Rabbit α-Syn1 | 5% Milk | 1:100,000 | Millipore AB1543<br>(Etobicoke, Ontario, Canada) |
| | Guinea Pig α-vGLUT1 | 5% Milk | 1:5,000 | Synaptic Systems 135304<br>(Goettingen, Germany) |
| | Rabbit α-PSD95 | 5% Milk | 1:5,000 | Cell Signaling Technology 2507S<br>(Danvers, MA, USA) |
| | Mouse α-GAD67 | 5% Milk | 1:10,000 | Advanced ImmunoChemical Inc<br>(Long Beach, CA, USA) |
| | Rabbit α-GPHN | 5% Milk | 1:2,000 | Cell Signaling Technology 14304S |
| 2° | Goat α-Rabbit HRP | - | 1:10,000 | Vector Labs PI-1000 (Burlingame,<br>CA, USA) |
| | Goat α-GuineaPig HRP | - | 1:10,000 | Invitrogen A18769<br>(Massachusetts, USA) |
| | Horse α-Mouse HRP | - | 1:10,000 | VectorLabs (Burlingame, CA, USA) |

**Supplementary Table 2:**
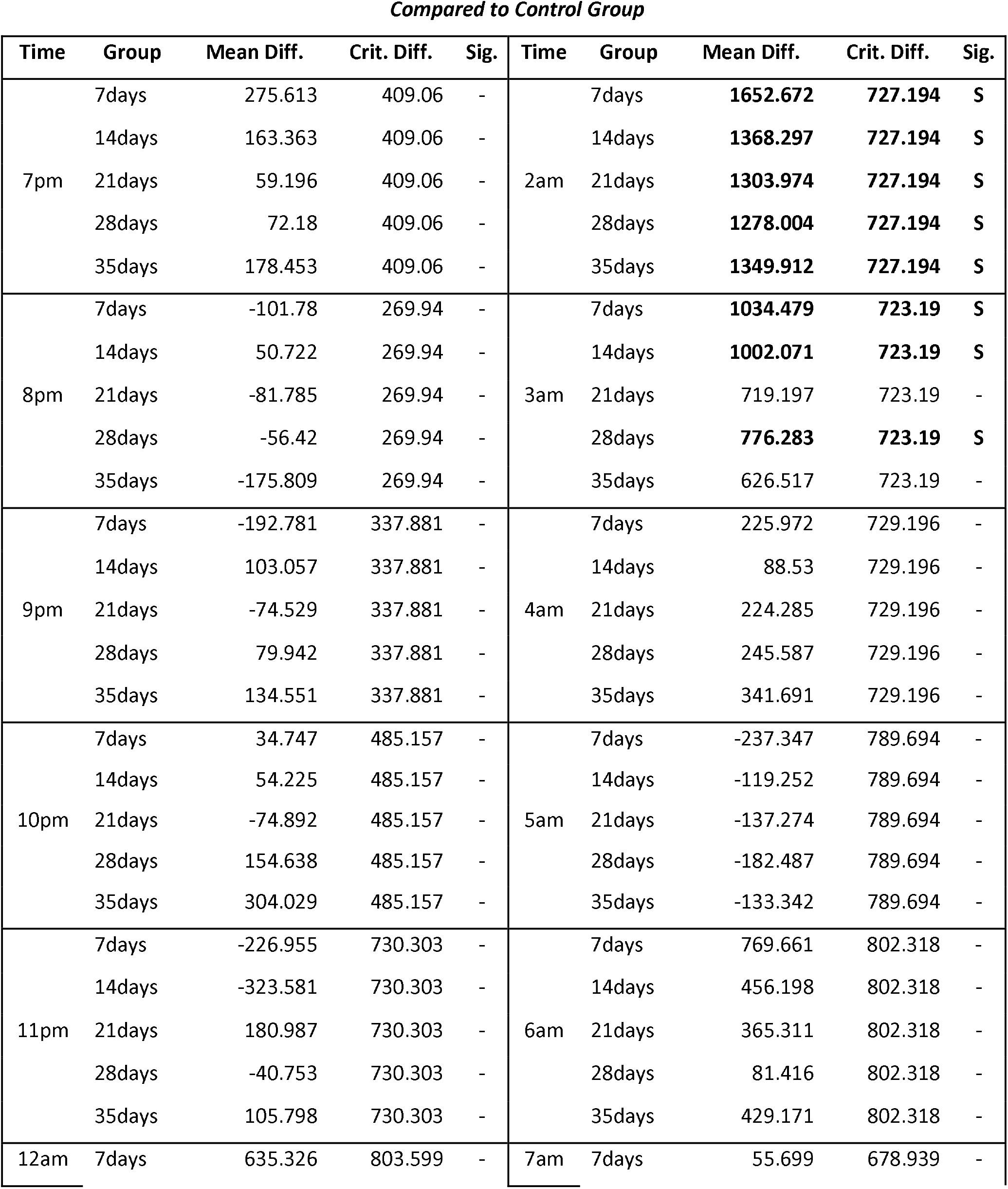
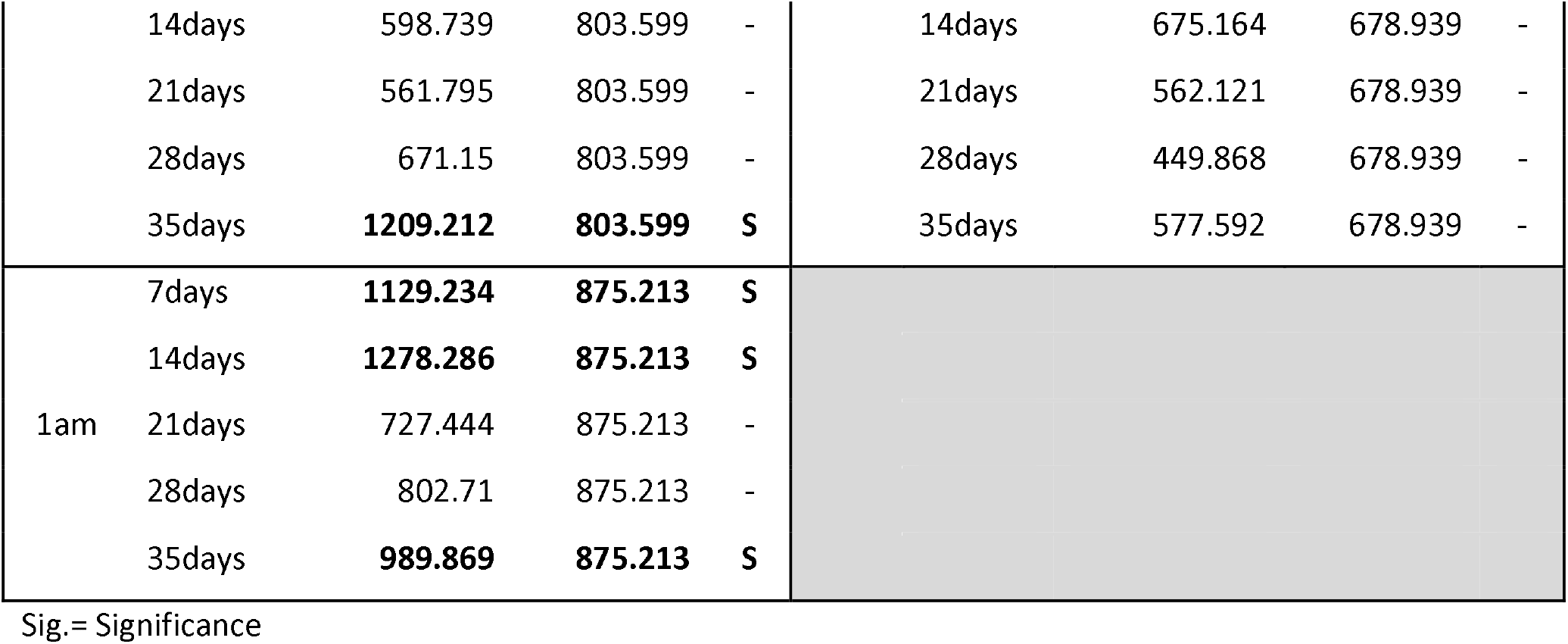
Dunnet’s Significance Table - Shelter Zone

**Supplementary Table 3:** Marker x Behavior Correlation Table. Underlined p-values remained significant after FDR correction (Benjamini-Hochberg).

| OVERALL |  | Pearson's Regression |  |  |  |  |
| --- | --- | --- | --- | --- | --- | --- |
|  |  | Weight Gain | RA FZ | RA Shelter | Sucrose | Z-Score |
| GLIA | GLT1 | 0.31 (0.1) | 0.74 (-0.18) | 0.57 (-0.22) | <u>0.0001</u> (0.4) | <u>0.004</u> (-0.3) |
|  | GFAP | 0.37 (-0.09) | 0.81 (-0.02) | 0.54 (-0.06) | <u>0.003</u> (0.3) | 0.13 (-0.15) |
|  | GS | 0.62 (-0.05) | 0.89 (0.01) | <u>0.07</u> (-0.19) | <u>0.07</u> (0.19) | 0.19 (-0.13) |
| Synaptic | vGLUT | 0.18 (0.13) | 0.2 (-0.1) | 0.85 (0.03) | 0.83 (0.05) | <u>0.008</u> (-0.27) |
|  | Syn1 | 0.42 (0.08) | <u>0.02</u> (-0.23) | <u>0.02</u> (-0.23) | <u>0.008</u> (0.26) | <u>0.0015</u> (-0.32) |
|  | PSD95 | 0.87 (0.01) | 0.61 (-0.05) | 0.23 (-0.12) | 0.11 (0.17) | <u>0.08</u> (-0.18) |
| GABA | GAD67 | 0.42 (0.08) | 0.28 (-0.1) | 0.43 (-0.12) | 0.48 (0.07) | <u>0.0026</u> (-0.3) |
|  | GPHN | 0.35 (0.1) | <u>0.05</u> (0.2) | <u>0.002</u> (0.3) | 0.93 (0.01) | 0.65 (-0.04) |
|  | SST | 0.13 (-0.15) | 0.41 (-0.02) | 0.25 (-0.1) | <u>0.09</u> (0.17) | 0.39 (-0.09) |
|  | PV | 0.6 (0.05) | 0.98 (0.00) | 0.8 (-0.02) | 0.91 (0.00) | 0.71 (-0.04) |
|  | VIP | <u>0.018</u> (0.24) | 0.81 (0.02) | 0.78 (0.03) | 0.49 (0.06) | 0.79 (0.02) |

| OVERALL |  | Spearman's Regression |
| --- | --- | --- |
|  |  | Coat State |
| GLIA | GLT1 | 0.39 (-0.09) |
|  | GFAP | 0.88 (-0.01) |
|  | GS | 0.96 (0.04) |
| Synaptic | vGLUT | <u>0.001</u> (-0.33) |
|  | Syn1 | <u>0.099</u> (-0.23) |
|  | PSD95 | 0.23 (-0.13) |
| GABA | GAD67 | <u>0.0003</u> (-0.4) |
|  | GPHN | <u>0.03</u> (-0.22) |
|  | SST | 0.41 (0.084) |
|  | PV | 0.61 (0.05) |
|  | VIP | 0.14 (0.15) |

$p(Rho)$
| MALE |  | Weight Gain | RA FZ | RA Shelter | Sucrose | Z-Score |
| --- | --- | --- | --- | --- | --- | --- |
| GLIA | GLT1 | <u>0.04</u> (0.29) | <u>0.09</u> (-0.24) | <u>0.003</u> (-0.5) | <u>0.003</u> (0.4) | <u>0.0004</u> (-0.4) |
|  | GFAP | <u>0.008</u> (0.39) | 0.27 (-0.16) | <u>0.004</u> (-0.46) | <u>0.0003</u> (0.5) | <u>0.04</u> (-0.29) |
|  | GS | <u>0.09</u> (0.25) | 0.33 (-0.1) | <u>0.01</u> (-0.34) | <u>0.007</u> (0.4) | <u>0.07</u> (-0.26) |
| Synaptic | vGLUT | 0.12 (0.23) | 0.83 (-0.03) | 0.9 (0.01) | 0.55 (-0.09) | 0.25 (-0.17) |
|  | Syn1 | 0.75 (0.08) | <u>0.01</u> (-0.35) | <u>0.03</u> (-0.3) | <u>0.02</u> (0.33) | <u>0.01</u> (-0.36) |
|  | PSD95 | 0.5 (0.1) | 0.18 (-0.2) | 0.37 (-0.13) | <u>0.02</u> (0.32) | <u>0.06</u> (-0.18) |
| GABA | GAD67 | 0.97 (0.004) | 0.25 (-0.1) | 0.12 (-0.2) | <u>0.09</u> (0.25) | <u>0.01</u> (-0.3) |
|  | GPHN | 0.24 (0.17) | 0.16 (0.2) | <u>0.09</u> (0.2) | 0.68 (-0.06) | 0.79 (-0.04) |
|  | SST | <u>0.01</u> (0.35) | 0.39 (-0.12) | <u>0.026</u> (-0.33) | <u>0.07</u> (0.26) | 0.25 (-0.17) |
|  | PV | 0.9 (0.02) | 0.77 (-0.04) | 0.29 (-0.16) | 0.91 (0.02) | 0.96 (-0.007) |
|  | VIP | 0.17 (0.2) | 0.84 (0.03) | 0.21 (0.18) | 0.63 (0.07) | 0.82 (0.03) |

| MALE |  | Coat State |
| --- | --- | --- |
| GLIA | GLT1 | <u>0.08</u> (-0.25) |
|  | GFAP | 0.63 (0.07) |
|  | GS | 0.64 (0.07) |
| Synaptic | vGLUT | <u>0.02</u> (-0.34) |
|  | Syn1 | 0.23 (-0.17) |
|  | PSD95 | 0.25 (-0.17) |
| GABA | GAD67 | <u>0.01</u> (-0.38) |
|  | GPHN | 0.18 (-0.19) |
|  | SST | 0.72 (0.05) |
|  | PV | 0.93 (0.01) |
|  | VIP | 0.92 (0.01) |

| FEMALE |  | Weight Gain | RA FZ | RA Shelter | Sucrose | Z-Score |
| --- | --- | --- | --- | --- | --- | --- |
| GLIA | GLT1 | 0.84 (0.03) | 0.4 (-0.12) | 0.85 (0.02) | <u>0.006</u> (0.4) | 0.45 (-0.1) |
|  | GFAP | 0.45 (0.11) | 0.68 (-0.06) | 0.63 (0.07) | 0.16 (0.2) | 0.15 (0.2) |
|  | GS | 0.3 (0.15) | 0.14 (0.21) | 0.75 (0.04) | 0.44 (-0.1) | 0.65 (0.06) |
| Synaptic | vGLUT | 0.75 (0.05) | 0.16 (-0.2) | 0.6 (0.08) | 0.1 (0.24) | <u>0.005</u> (-0.4) |
|  | Syn1 | 0.33 (0.14) | 0.72 (-0.05) | 0.69 (-0.05) | 0.43 (0.11) | 0.41 (-0.12) |
|  | PSD95 | 0.74 (0.05) | 0.77 (0.04) | 0.32 (-0.14) | 0.85 (0.02) | 0.41 (-0.12) |
| GABA | GAD67 | 0.33 (0.14) | 0.86 (-0.02) | 0.88 (0.02) | 0.48 (-0.1) | 0.1 (-0.23) |
|  | GPHN | 0.93 (0.002) | 0.62 (0.07) | <u>0.09</u> (0.24) | 0.12 (0.22) | <u>0.08</u> (-0.25) |
|  | SST | 0.87 (0.02) | 0.37 (-0.13) | 0.9 (0.01) | 0.46 (0.17) | 0.55 (-0.09) |
|  | PV | 0.5 (0.1) | 0.45 (-0.1) | 0.47 (-0.1) | 0.43 (0.12) | 0.12 (-0.22) |
|  | VIP | <u>0.057</u> (0.28) | 0.36 (-0.13) | 0.22 (-0.1) | 0.36 (0.13) | 0.67 (0.06) |

| FEMALE |  | Coat State |
| --- | --- | --- |
| GLIA | GLT1 | 0.47 (0.1) |
|  | GFAP | 0.26 (-0.16) |
|  | GS | 0.99 (-0.002) |
| Synaptic | vGLUT | <u>0.058</u> (-0.27) |
|  | Syn1 | 0.72 (-0.05) |
|  | PSD95 | 0.72 (-0.05) |
| GABA | GAD67 | <u>0.007</u> (-0.4) |
|  | GPHN | <u>0.017</u> (-0.5) |
|  | SST | 0.74 (0.05) |
|  | PV | 0.52 (-0.09) |
|  | VIP | 0.27 (0.16) |

$p(Rho)$

**Supplementary Table 4:**
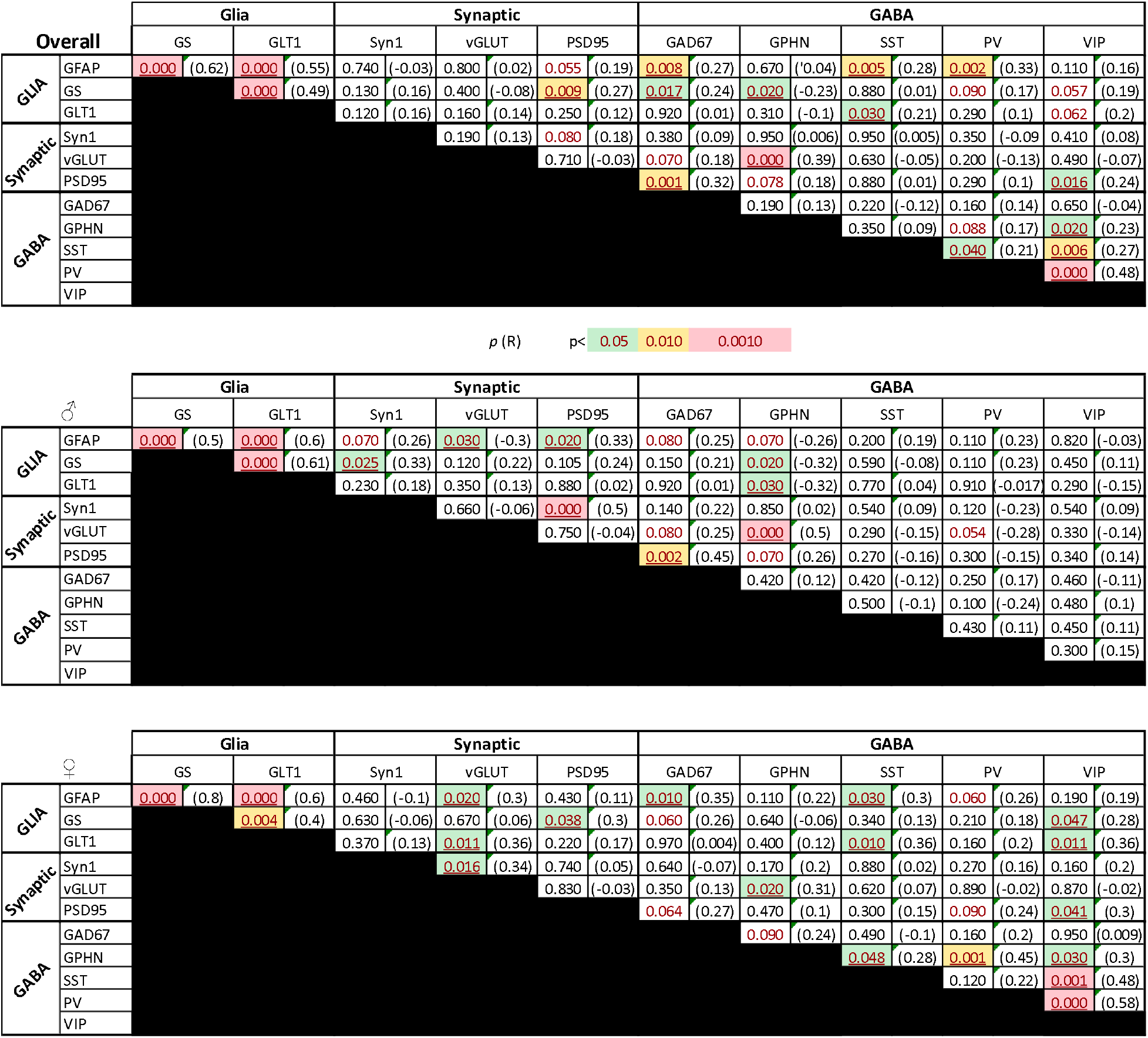
Marker x Marker Correlation Table. Underlined p-values remained significant after FDR correction (Benjamini-Hochberg).

### - Supplementary Figures -

**Supplementary Figure 1:**
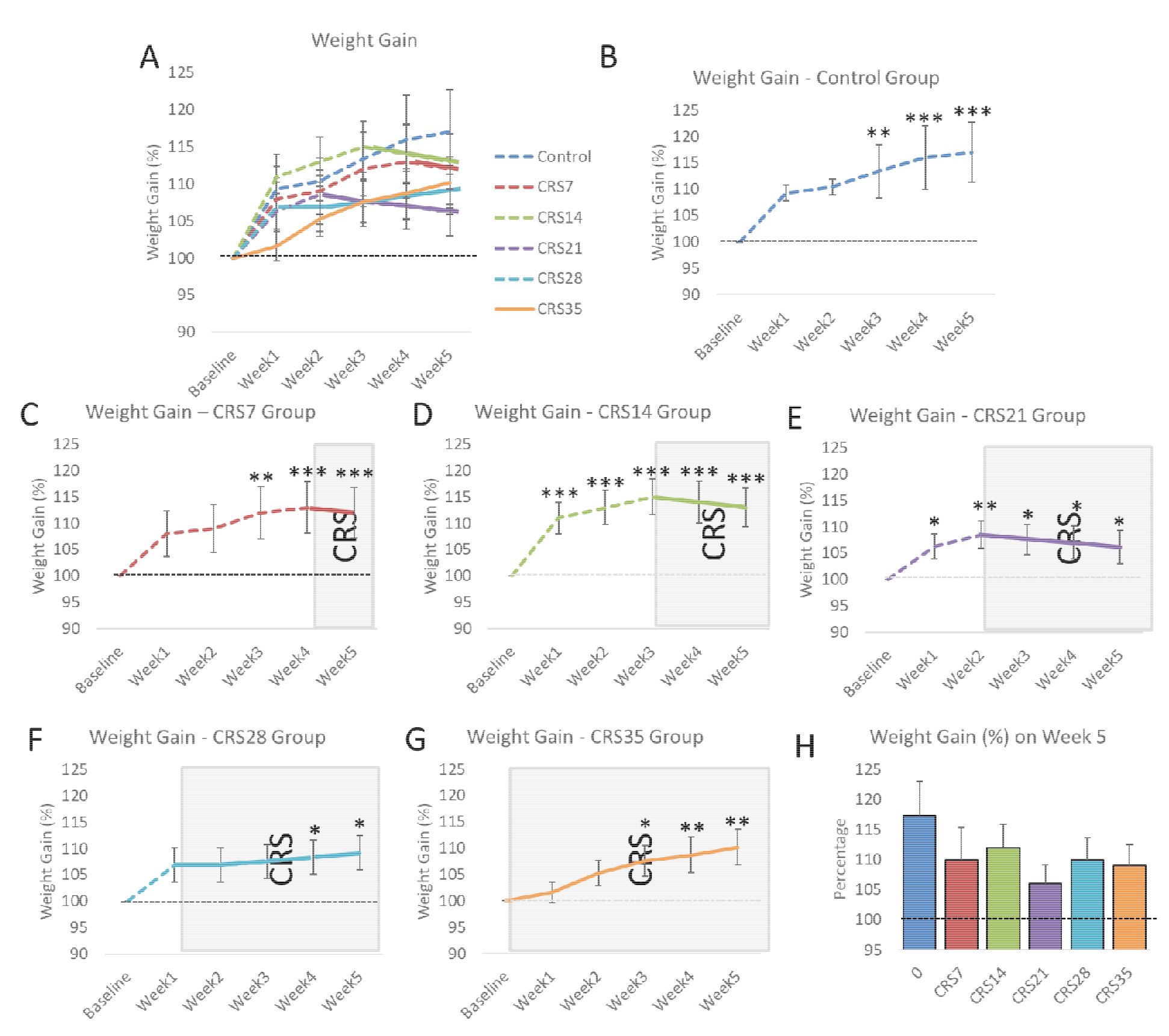
Evolution of weight gain over weeks. Mice were weighted on a weekly basis over the course of the experiment. Weight gain was measured from their baseline weight (representing 100%), and was calculated every week. Overall weight gain results including all groups is presented in the panel A, and split per group in the panels B through G. The black dotted line represents 100%. The colored dotted lines represent the weight gain when the mice are not subjected to CRS. Plain lines represent the period when the are subjected to CRS, also presented by the gray zone in the background. Panel H presents the overal weight gain measured after Week 5, i.e after completion of the entire study. *p<0.05;**p<0.01, ***p<0.001 compared to Baseline.

**Supplementary Figure 2:**
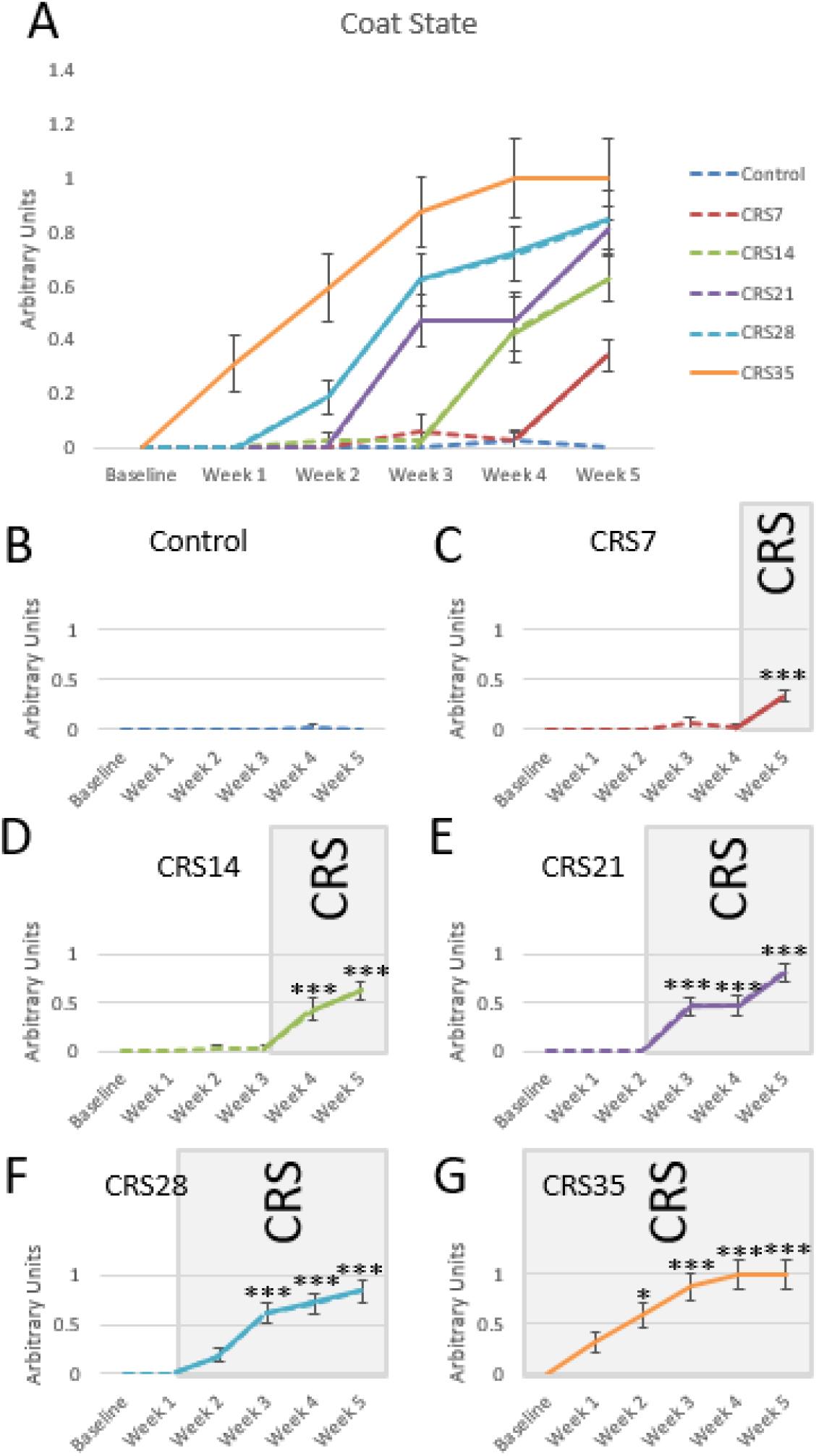
Evolution of Coat State over the weeks. Coat state was assessed every week, based on the method described in Yalcin et al (2015). Panel A provided an overview of the coat state in all groups, throughout the entire study. Panels B through G represent the different groups starting with the Control group (B) and finishing with the CRS35 group (G). The colored dotted lines represent the coat state when the mice are not subjected to CRS. Plain lines represent the period when the are subjected to CRS, also presented by the gray zone in the background. *p<0.05;**p<0.01, ***p<0.001 compared to Baseline.

**Supplementary Figure 3:**
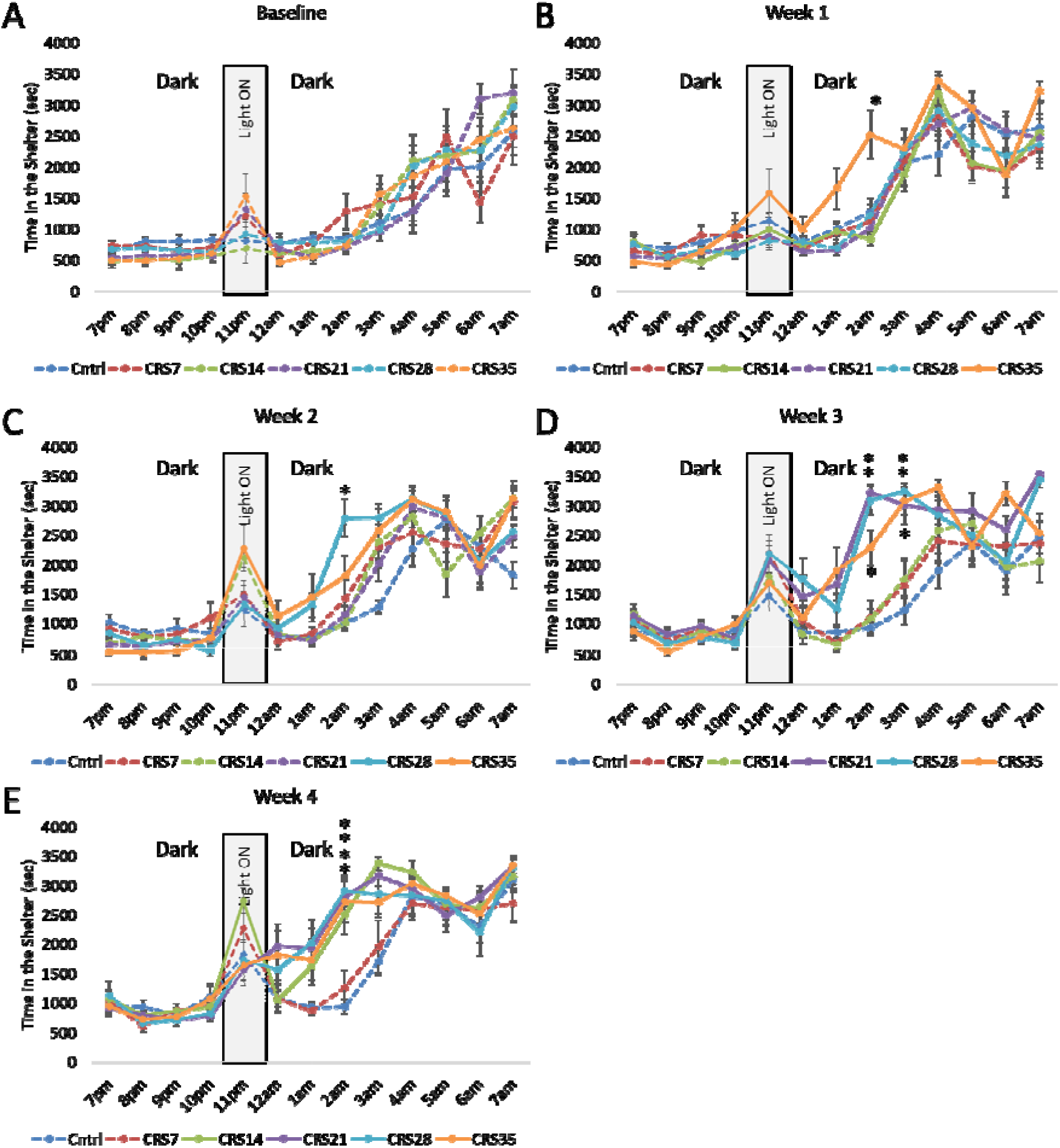
Assessment of anxiety-like behaviors in the Phenotyper on a weekly basis. Mice from all groups were tested in the Phenotyper test, whether their exposure to CRS had started or not. Before initiation of the CRS paradigm, mice were tested in the Phenotyper test for baseline activity (A), showing no difference between groups, assigned artificially at this stage. Then, on Week 1 (B), only the animals from the CRS35 group had started CRS, while the others were kept as control. On this week, we showed a significant effect of CRS, characterized by increasing time in the shelter, in particular at 2am. This is observed on Week 2 (C), 3 (D) and 4 (E). In this figure, groups subjected to CRS are represented with plain lines. Groups that were not subjected to CRS yet, are presented with dotted lines. The gray zone in the background of each graph represents the time when the light was turned ON, as an acute challenge. Data are presented as average per group ± SEM. *p<0.05 compared to Control mice.

**Supplementary Figure 4:**
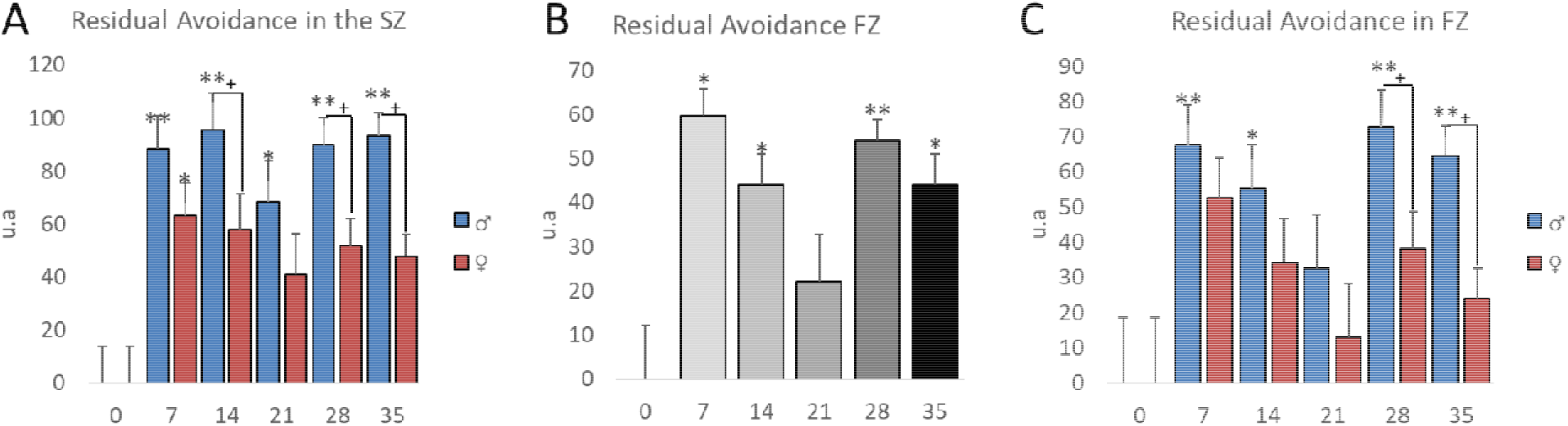
Residual avoidance in the Shelter and in the Food Zones, including sex differences. Residual avoidance was calculated in a sex-dependent manner, with male and female RA from the Control group both being equal to 0. Overall effect of CRS Duration was significant, even after splitting the dataset by sex, in the Shelter Zone (SZ; Panel A). The effect of sex showed that males from the CRS14, CRS28 and CRS 35 groups exhibited an overall higher RA score than females. Residual avoidance was calculated in the Food Zone (B) and showed significant effect of the CRS Duration (F_(5;82)_=6.9; p<0.001), an effect of Sex (F_(1;82)_=9.5; p=0.002) and no Duration*Sex interaction (p>0.05). *Post hoc* Dunnett’s test revealed an increase in RA score compared to Control after 7, 14, 28 and 35 days of CRS (p<0.05), but not after 21 days of CRS (p=0.6). In the FZ, the effect of sex was characterized and showed higher scores in males after 28 and 35 days of CRS (C). *p<0.05;**p<0.01 compared to Control; +p<0.05 compared to Females.

**Supplementary Figure 5:**
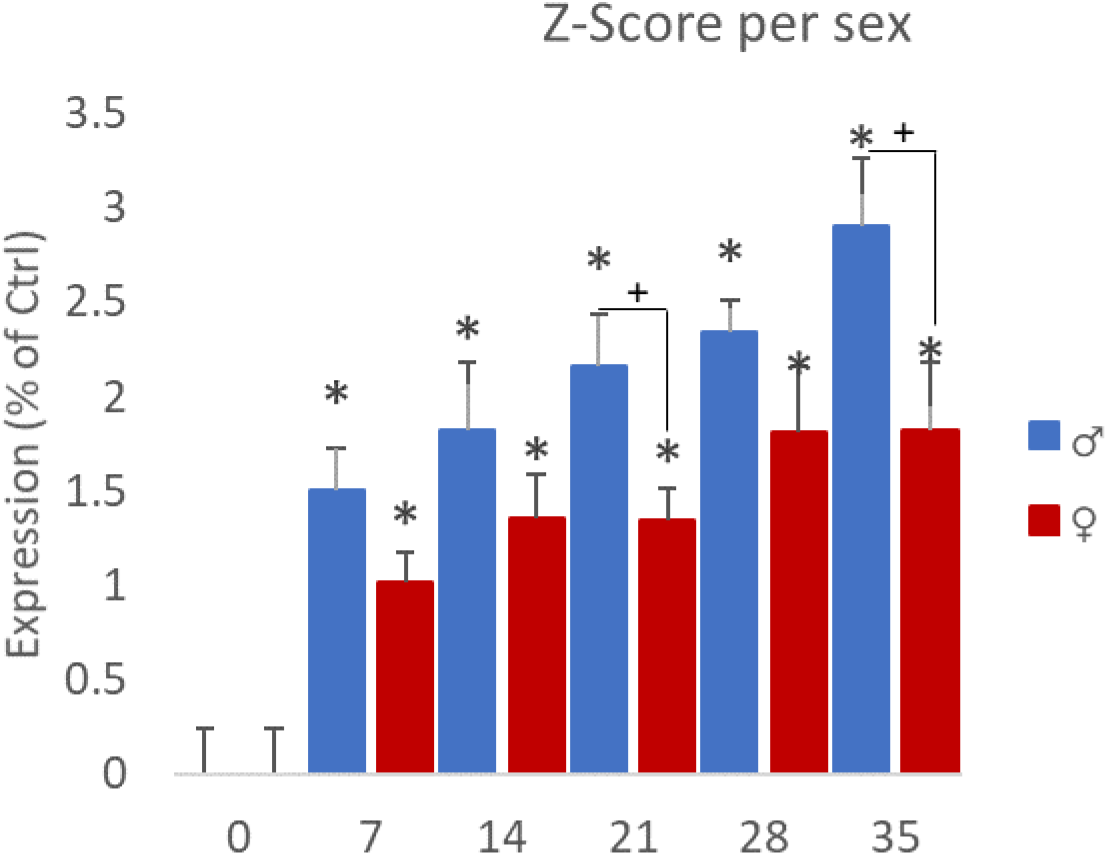
Z-Score per sex. Z-scores were calculated in a sex-dependent manner, with both z-scores from male and female control groups being equal to 0. ANOVA performed on the Z-Score showed a significant effect of CRS Duration (F_(5,82)_=18.2, p<0.001), a significant effect of Sex (F_(1,82)_=12.9, p=0.0006) and no CRS Duration*Sex interaction (p>0.5). *Post hoc* analyses show that in both males and females there is a significant increase of the z-score in all CRS duration groups, compared to the Control group (ps<0.01). The effect of Sex is explained by a Z-score higher in males than females. *p<0.05 compared to Control; +p<0.05 compared to Females.

**Supplementary Figure 6:**
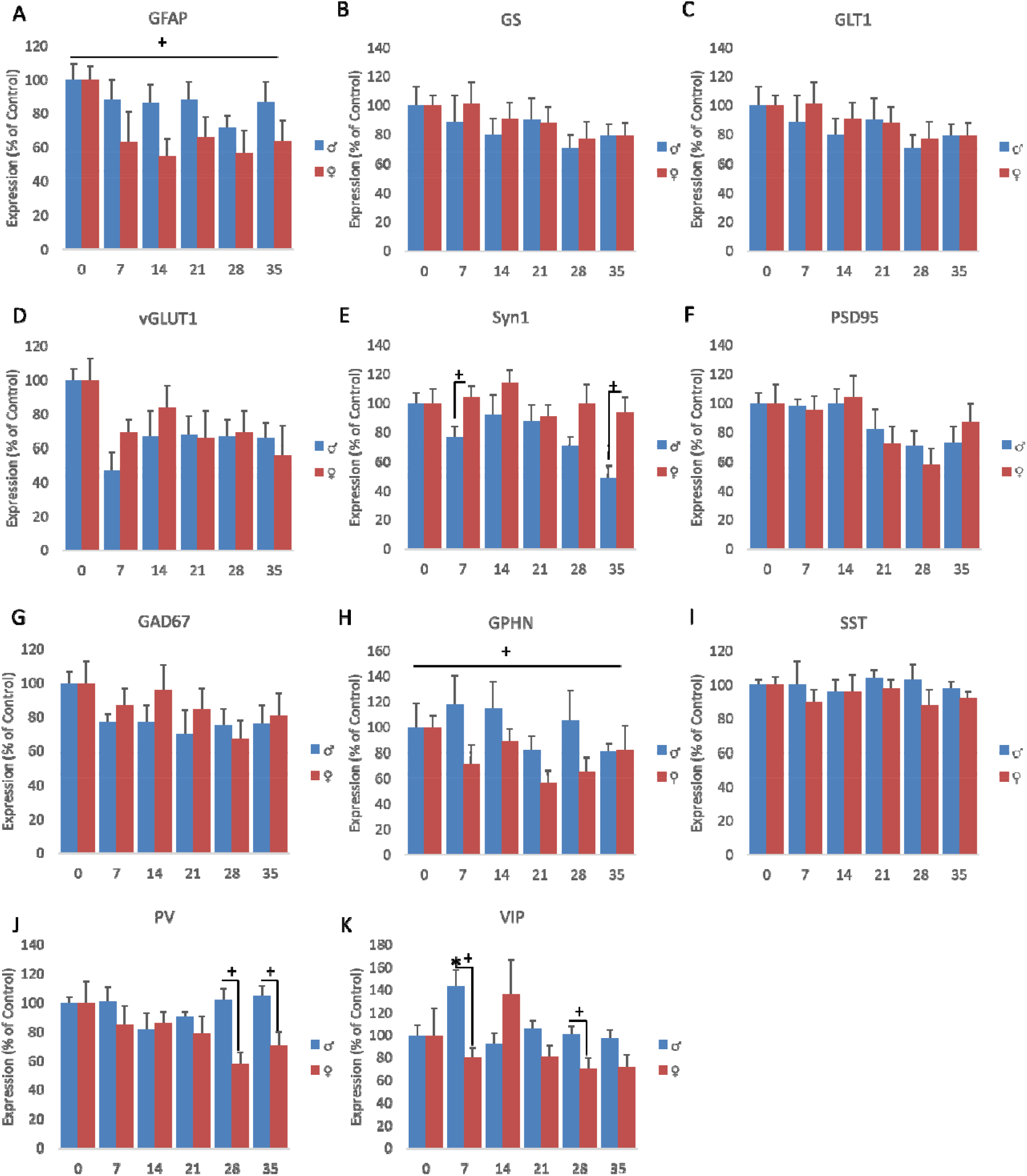
Cellular markers per sex. Expression levels of each marker was assessed in a sex-dependent manner. Statistical analyses performed on GFAP expression levels showed an overall effect of sex (A). No effects of sex were observed on expression levels of GS (B), GLT1 (C) or vGLUT1(D). Syn1 expression levels were significantly lower in males compared to females in the CRS7 and CRS35 groups (E). No effects of sex were observed on expression levels of PSD95 (F) and GAD67 (G). An overall effect of sex was identified on GPHN expression levels (H). No effects of sex were observed on expression levels of SST (I). A significant effect of sex was identified in PV expression levels, characterized by lower expression levels in females compared to males in CRS28 and CRS35 groups (J). Finally, an interaction between CRS Duration*Sex was observed on VIP expression levels, which was characterized by higher expression levels in male compared to females in the CRS7 and CRS28 groups (K). *p<0.05 compared to Control; +p<0.05 compared to Females.

**Supplementary Figure 7:**
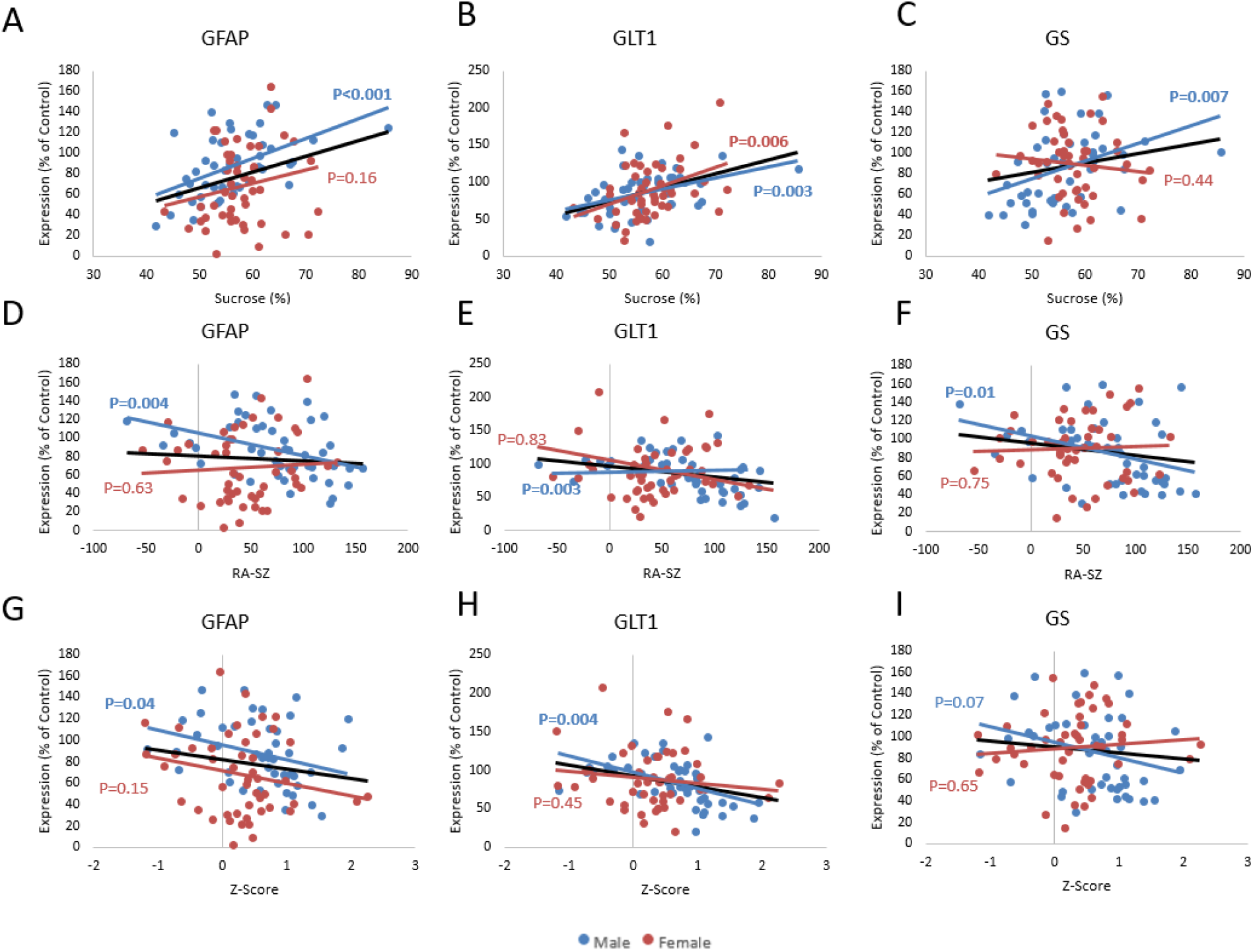
Behavioral outcomes and glial marker correlations split by sex. Pearson’s regression analyses were performed between glial markers and behavioral outcomes, and split by sex. GFAP, GLT1 and GS expression levels correlated with Sucrose consumption in male but not in females (A-C). GFAP, GLT1 and GS expression levels also correlated with residual avoidance in the shelter in males but not in females (D-F). Finally, GFAP and GLT1 expression levels in males, but not in females, correlated with z-scores (G-H), while GS expression levels were only trending towards significance (I).

**Supplementary Figure 8:**
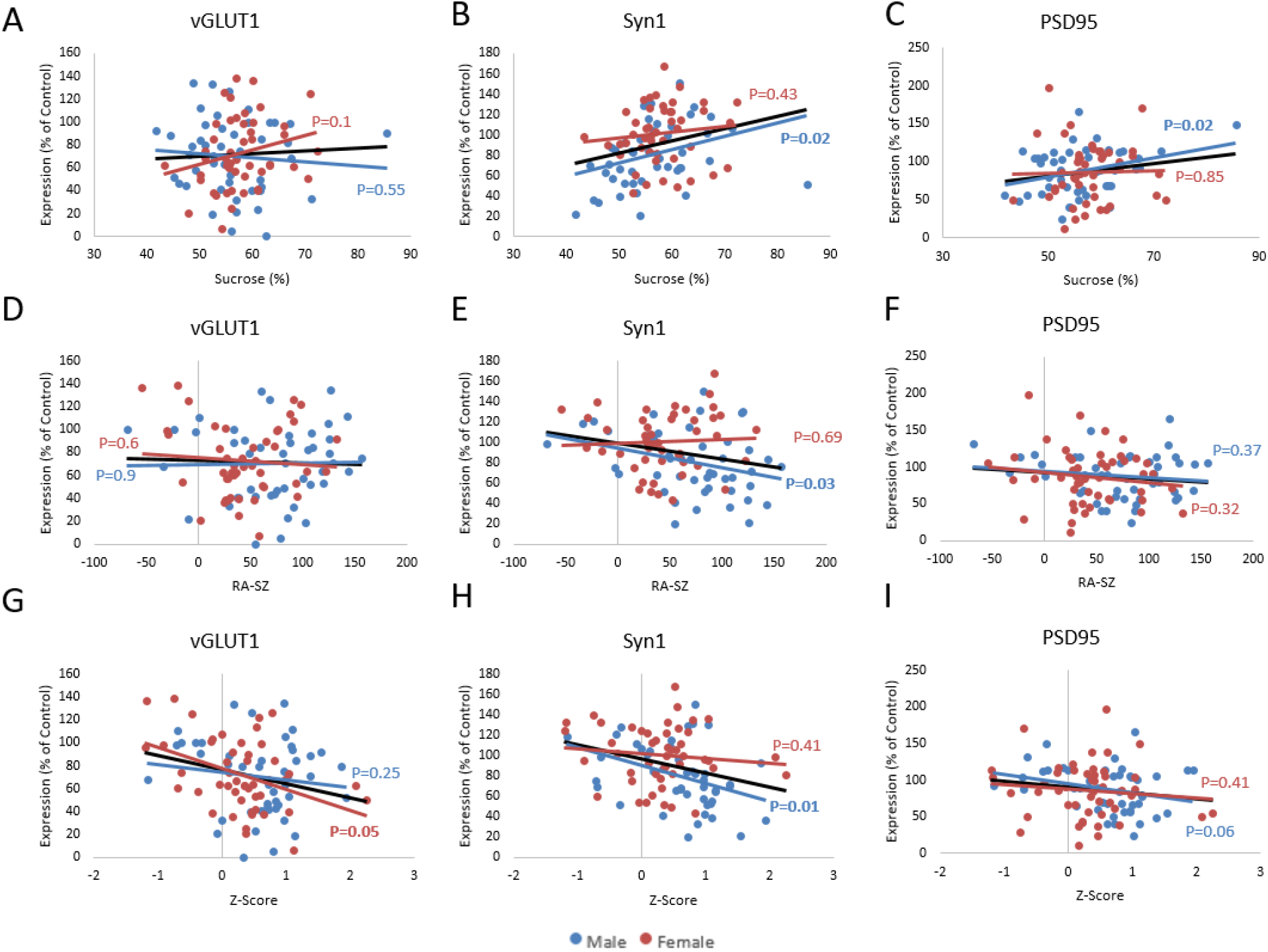
Behavioral outcomes and synaptic marker correlations split by sex. Pearson’s regression analyses were performed between synaptic markers and behavioral outcomes, and split by sex. vGLUT1 expression levels did not correlate with sucrose consumption, neither in males nor in females (A). In males, Syn1 and PSD95 expression levels correlated with sucrose consumption but not in females (B-C). Regarding correlating with residual avoidance in the shelter, vGLUT and PSD95 expression levels did not correlated with RA-SZ neither in males nor in females (D and F), while Syn1 expression levels correlated with RA-SZ in males only (E). vGLUT1 expression levels correlated with z-score in females but not in males (G). Syn1 expression levels correlated with z-score only in males (H). Finally, PSD95 expression levels did not correlate with z-scores, however levels in males were trending towards significance (I).

**Supplementary Figure 9:**
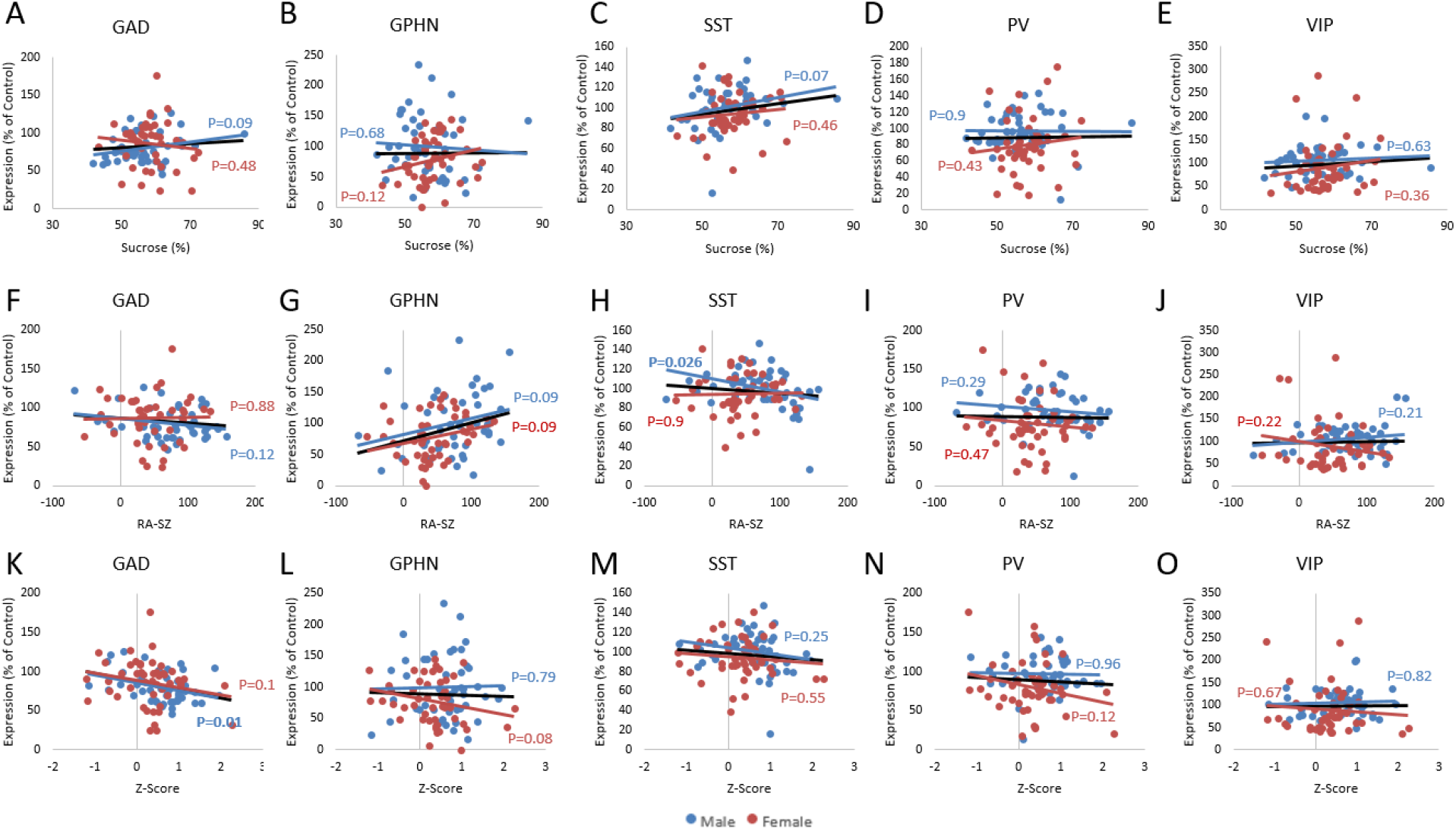
Behavioral outcomes and GABAergic marker correlations split by sex. Pearson’s regression analyses were performed between GABAergic markers and behavioral outcomes, and split by sex. Sucrose consumption did not correlate with expression levels of GAD (A), GPHN (B), SST (C), PV (D) or VIP (E) neither in males nor in females. Regarding residual avoidance in the shelter (RA-SZ), it did not correlate with GAD expression levels, neither in males nor in females (F). Expression levels of GPHN were trending in both males and females (G). RA-SZ correlated negatively with SST expression levels in males but not in females (H). Regarding PV and VIP expression levels, they did not correlate with RA-SZ, neither in males nor in females (I-J). Z-scores did not correlate with GAD expression levels (K), GPHN (L), SST (M), PV (N) or VIP (O), neither in males nor in females.

**Supplementary Figure 10.**
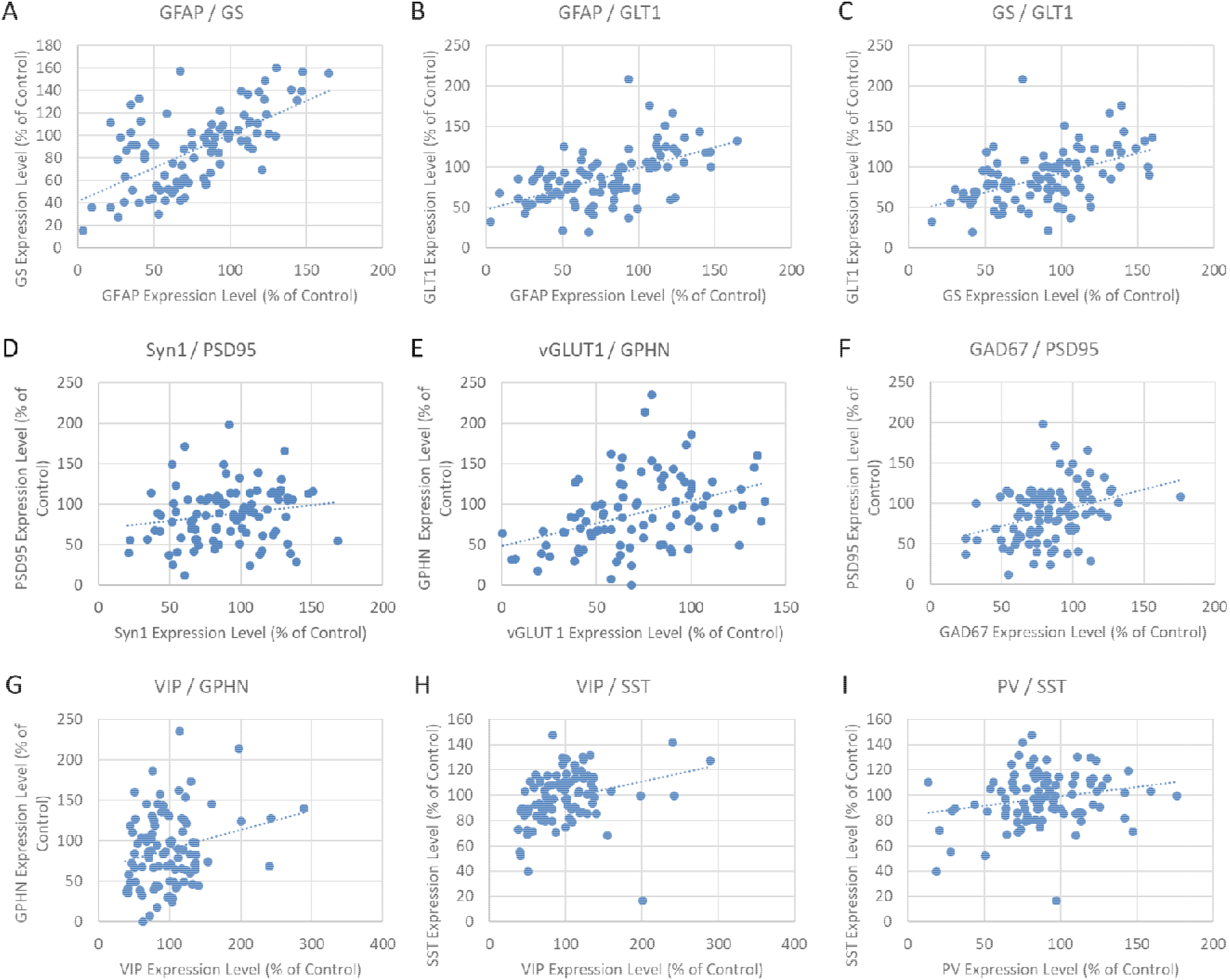
Between Markers Correlations. Pearson’s regression analyses between GFAP and GS (A), and GLT1 (B) showed significant positive correlation (ps<0.001). GS and GLT1 expression levels also correlated strongly with each other (p<0.001; C). Pearson’s regression analyses also showed a trend toward a positive correlation between PSD95 and Syn1 levels (R=0.18, p=0.08, D). vGLUT correlated with GPHN (p<0.001, E). PSD95 correlated with GAD67 (p=0.001,F). Finally, many of the GABAergic markers significantly correlated with each other. VIP expression levels correlated with GPHN (p=0.02, G) and SST (p=0.006, H). PV was correlated with SST (p=0.04, I).

